# TRPV2 interaction with small molecules and lipids revealed by cryo-EM

**DOI:** 10.1101/2020.08.10.242008

**Authors:** Anna D. Protopopova, Ruth A. Pumroy, Jeanne de la Roche, Ferdinand M. Haug, Bárbara B. Sousa, Pamela N. Gallo, Gonçalo J.L. Bernardes, Andreas Leffler, Vera Y. Moiseenkova-Bell

## Abstract

Transient receptor potential vanilloid 2 (TRPV2) plays a critical role in a variety of physiological and pathophysiological processes, putting TRPV2 on the list of important drug targets. Yet, specific TRPV2 agonists and antagonists are currently unavailable. Their development requires a precise knowledge of how the currently known non-specific small molecules interact with TRPV2 at the molecular level. Here we present TRPV2 structures in ligand-bound states resolved by cryo-electron microscopy in the presence of 2-aminoethoxydiphenyl borate (2-ABP), 2-APB with doxorubicin (DOXO), and ruthenium red (RR). We identified a novel 2-APB drug binding site between the S5 helix and S4-S5 linker on two adjacent TRPV2 monomers and determined the mechanism of TRPV2 pore block by RR. We also showed that a large organic molecule like DOXO can enter the TRPV2 pore in the presence of 2-APB. Additionally, we discovered a structural lipid bound in a unique position in the “vanilloid pocket”, which is absent in the 2-APB-bound state of the channel, allowing us to propose a model for TRPV2 channel gating. Together, this work provides a further understanding of TRPV2 function and a structural framework for the development of TRPV2-specific modulators.

## Introduction

Transient receptor potential vanilloid 2 (TRPV2) is a non-selective cation channel belonging to the vanilloid subfamily of the transient receptor potential (TRP) channels^1,2^. TRPV2 is expressed in a wide variety of human tissues, including the heart, white blood cells, the pancreas, the central and peripheral nervous systems, the bladder, and the prostate^3^. TRPV2 has been shown to play a significant role in maintaining physiological cardiomyocyte function, with a potential therapeutic use of TRPV2 blockade in cardiomyopathy^4–6^. TRPV2 is required for phagocytosis in macrophages^7,8^ and has also been shown to be crucial for insulin secretion in pancreatic β-cells^9,10^. Beyond its routine functions in healthy cells, TRPV2 has been shown to play a significant role in different forms of cancer^11^. It is downregulated in gliomas^12^, while activation of TRPV2 with cannabidiol (CBD) in glioblastoma cells facilitated cell death by increasing the uptake of cytotoxic chemotherapeutic drugs doxorubicine (DOXO), temozolomide, and carmustine^13^. Overexpression of TRPV2 mRNA and protein has been observed in bladder and prostate cancer, suggesting a potential role of TRPV2 in drug resistance and metastasis^14,15^. TRPV2’s role in both healthy tissues and in multiple forms of cancer makes it an interesting target for drug discovery and development of novel therapeutics in cancer, cardiovascular diseases, and additional pathophysiological conditions.

A key limiting factor in TRPV2 studies is the lack of specific modulators of the channel. There are several known non-specific exogenous activators of TRPV2, including 2-aminoethoxydiphenyl borate (2-APB), CBD and probenecid^2^. However, 2-APB has also been shown to activate TRPV1-3^16^ and inhibit TRPV6^17^, while CBD activates TRPV1-4^18^ and both ligands affect other ion channels and receptors^19–22^. While probenecid has been used in some TRPV2 studies as a channel activator^2^, it is not a commonly used modulator of channel activity *in vitro* and *in vivo*. Even less is known about specific TRPV2 inhibitors. The most effective known inhibitors are ruthenium red (RR) and trivalent cations, but these function as non-specific channel blockers^2,23,24^. Tranilast, an antiallergic drug, has been suggested to inhibit TRPV2^2^, but this was called into question when a recent study showed that tranilast has no direct effect on TRPV2 *in vivo*^25^.

To aid the rational development of modulators with increased efficacy and TRPV2 specificity, reliable structural data is required on known small molecule binding to TRPV2. Although structures of CBD-bound full-length wild-type (WT) rat TRPV2 in nanodiscs recently revealed the CBD binding site in a pocket located between the S5 and S6 helices of adjacent TRPV2 monomers^26^, all other information on potential TRPV2 ligand binding sites comes from other members of the TRPV family.

Vanilloids, like capsaicin and resiniferatoxin (RTX), activate only TRPV1 and bind to the pocket located above the S4-S5 linker between S3 and S4 of one monomer and S6 of an adjacent monomer^27–29^. This “vanilloid pocket” in TRPV2 diverges significantly from the TRPV1 pocket and has lost sensitivity to vanilloids, although it can be mutated to regain vanilloid sensitivity^30,31^ (**Figure S1A**). As in the TRPV1 apo structure, most TRPV structures^17,26,29,32–41^ have some lipid density in the vanilloid pocket. The proposed mechanism of ligand activation of TRPV1 is through displacing a lipid found in the vanilloid pocket, but it is unknown if this pocket plays a role in native TRPV2 activation.

2-APB can activate TRPV1-TRPV3^16^ and three TRPV3 structural studies have recently been published featuring 2-APB bound states^41–43^. This work has led to a consensus TRPV3 2-APB binding site near human His426 between the Pre-S1 helix and TRP helix, which agrees with previous functional data^44^. However, this critical residue is not conserved in TRPV1 and TRPV2^44^ (**Figure S1A**). Thus, the 2-APB binding site in TRPV3 is not applicable to TRPV2, and presents the possibility of a new, unique binding site that could be used to develop TRPV2-specific therapeutics.

Structural information on potential TRPV2 inhibition is absent. RR is a common cation channel pore blocker^45^, but so far there is limited structural information about how it interacts with cation channels^45,46^. Thus, TRPV2 is likely to reveal a new RR binding site that could be conserved in other TRPV channels.

Transport of large organic cations into the cell through the activated TRPV2 channel has been shown in multiple cell lines via the uptake of fluorescent dyes and drugs^13,47,48^. Additionally, several ion channels including P2X receptor channels^49–51^ and other TRP channels^52–54^ have also been found to permit large molecules into cells. This phenomenon was initially explained by a pore dilation mechanism^51,55^, but more recently it has been suggested that the normal conducting state is sufficient to permit large organic cations^56^. Therefore, structural information on this aspect of TRPV2 pharmacology could also lead to more specific therapeutic approaches.

To address these needs, we used cryo-electron microscopy (cryo-EM) to determine the structures of WT rat TRPV2 in the presence of 2-APB and RR in lipid membranes, which revealed a unique mechanism of TRPV2 modulation by 2-APB and a conceivably conserved mechanism of TRPV1-4 channel block by RR. We have identified a novel binding pocket for 2-APB between the S5 helix and the S4-S5 linker of two adjacent monomers and a RR binding site at the junction between the bottom of the pore helix and the pore loop of the same monomer. We also revealed that large organic molecules, like DOXO, can enter the pore of TRPV2 upon its activation by 2-APB, which is consistent with previous observations using cell-based assays for TRPV2 and other TRP channels. Furthermore, we observed a unique lipid in the TRPV2 vanilloid pocket, which may need to be displaced for 2-APB channel activation. Together, our results lay the foundation for uncovering modes of TRPV2 channel interaction with small molecules, channel gating, and for future development of TRPV2-specific modulators.

## Results

### Structure of TRPV2 in 2-APB-bound state

We recently reported the first structures of a WT rat TRPV2 channel in nanodiscs in two distinct apo states and two distinct CBD-bound states^26^. To further understand TRPV2 channel pharmacology and activation mechanisms, we compared the effect of CBD and 2-APB on TRPV2 gating using both single channel (**Figure S2A-D**) and whole-cell recordings (**Figure S2E**) on HEK293T cells transiently transfected with WT rat TRPV2. In inside-out recordings, 10 μM CBD (**Figure S2C-D**) evoked single channel opening with a three times lower absolute open probability as compared to 100 μM 2-APB (**Figure S2A-B**) (n=3). In wholecell recordings, 30 μM CBD evoked very slowly activating and inactivating inward currents which were markedly different from the rapidly activating and inactivating currents evoked by 1 mM 2-APB (n=5) (**Figure S2E**). These data not only suggest that 2-APB is a more potent activator than CBD, but also indicate distinct TRPV2 activation mechanisms by CBD and 2-APB.

We next obtained a 2-APB-bound structure of TRPV2 by cryo-EM in order to determine the 2-APB binding site. We incubated WT rat TRPV2 reconstituted in nanodiscs with 1 mM 2-APB for 30 minutes before preparing grids for cryo-EM. The resulting dataset yielded a single stable structure with C4 symmetry and an overall resolution of 3.6 Å (TRPV2_2APB_) (**Figure S3-S4, Table S1**). This structure revealed a putative 2-APB binding site between S5 of one TRPV2 monomer and the S4-S5 linker of the adjacent monomer (**Figure 1A-B, S5A-B**). The proposed TRPV2 2-APB binding site is located at the N-terminal end of the S4-S5 linker, where the two aromatic rings coordinate with His521 (**Figure 1B, S5B**). In the WT rat TRPV2 apo state 1 (TRPV2_APO_1_, PDB 6U84)^26^, His521 may be involved in stabilizing the interaction between the S4-S5 linker and S5 from an adjacent monomer, as it appears to form a network of cation-π interactions with Arg539 and Tyr525 (**Figure 1C, S5C**). 2-APB binding disrupts this network, interacting with His521 and potentially acting as a wedge to force apart the S4-S5 linker and S5 (**Figure 1B, S5B**). Notably, a histidine in the His521 position is conserved in TRPV2 across a wide range of mammals but is not found in other members of the TRPV family (**Figure S1B**).

**Figure 1.**
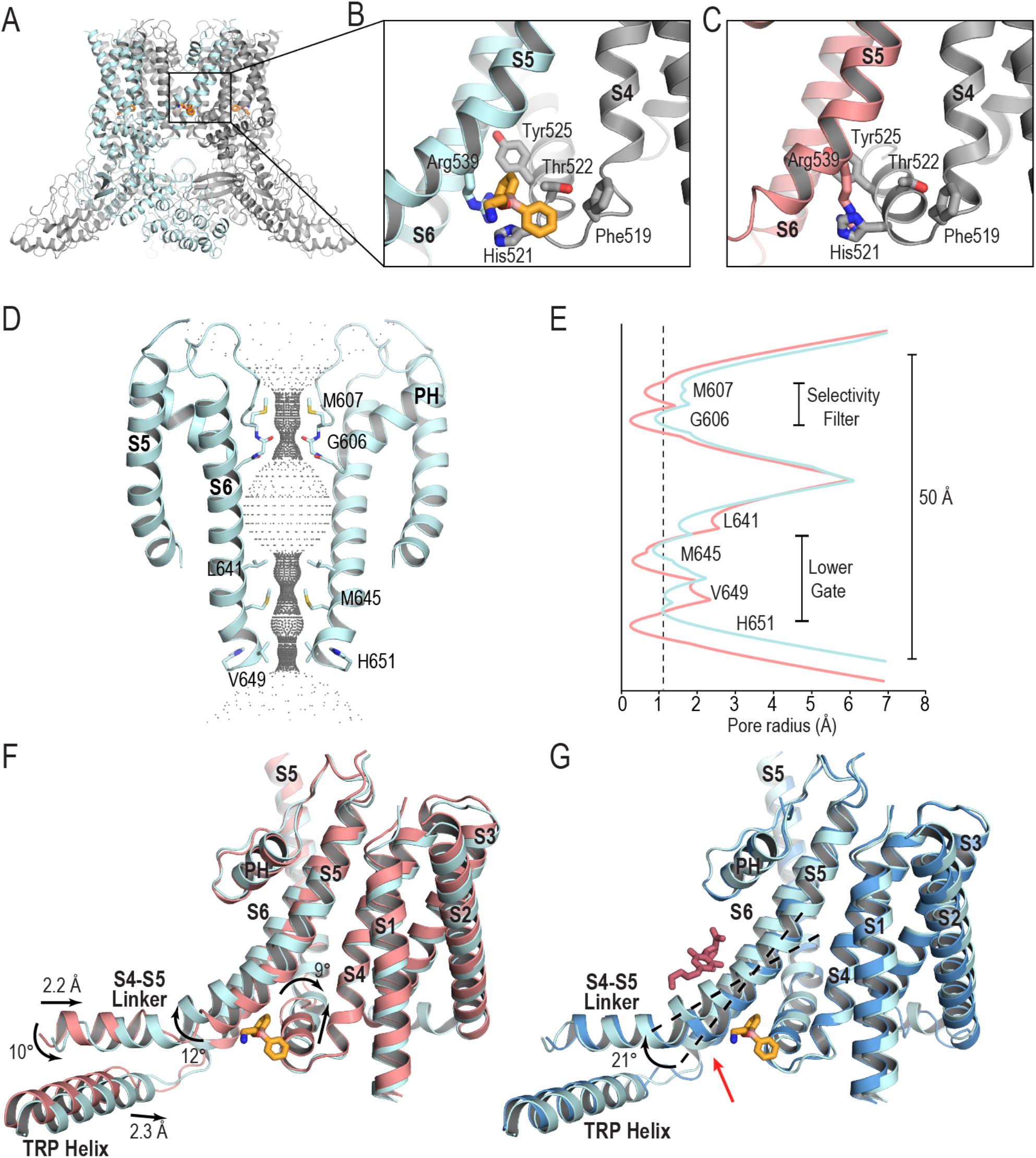
2-APB binds at the S4-S5 linker. (A) Cartoon representation of the TRPV2_2APB_ model. Detailed view of (B) the 2-APB binding site and (C) the same site from the TRPV2_APO_1_ model. One monomer of TRPV2 is colored grey, the adjacent monomer is colored pale cyan (TRPV2_2APB_) or salmon (TRPV2_APO_1_; PDB 6U84); 2-APB is depicted as orange sticks. D) Pore profile of TRPV2_2APB_ and (E) a graphical representation of the TRPV2_2APB_ (pale cyan) and TRPV2_APO_1_ (PDB 6U84) (salmon) pore profiles. Comparison of shifts between (F) TRPV2_2APB_ (pale cyan) and TRPV2_APO_1_ (PDB 6U84) (salmon) and (G) TRPV2_2APB_ (pale cyan) and TRPV2_CBD_2_ (PDB 6U88) (blue). Structures are aligned based on the tetrameric pore domain. 2-APB is depicted as orange sticks; CBD is depicted as pink sticks.

To validate this binding site for 2-APB, we used whole-cell patch recordings on HEK293T cells transiently transfected with either the WT rat TRPV2 or with three TRPV2 mutants around the proposed binding site: His521Ala, Thr522Ala, and Arg539Ala. While all mutants retained 2-APB-sensitivity, concentration-dependent activation by 2-APB was reduced in the His521Ala mutant compared to WT rat TRPV2. The mutants Arg539Ala and Thr522Ala displayed unchanged sensitivities to 2-APB (**Figure S6)**. However, the inward current densities evoked by 1mM 2-APB did not reveal any significant differences between WT rat TRPV2 and the mutant constructs (p=0.29, one-way ANOVA) (**Figure S6**). We also performed calcium-imaging experiments with HEK293T cells transiently transfected with WT human TRPV2 and confirmed significant stimulation (p<0.001; two-way ANOVA) of calcium influx evoked by 1 mM 2-APB (**Figure S7**). Cells transfected with an Arg537Ala human TRPV2 mutant, equivalent to the Arg539Ala rat TRPV2 mutant, showed the same level of stimulation as the WT channel. However, in cells transfected with a His519Ala human TRPV2 mutant, equivalent to the His521Ala rat TRPV2 mutant, the level of calcium influx evoked by 2-APB was diminished and similar to cells transfected with the pcDNA3.1(+) empty vector. These data support a potential interaction of 2-APB and TRPV2 via His519/521 for both the rat and human channels.

Although we have trapped the channel in a 2-APB-bound state, the TRPV2_2APB_ ion conducting pathway is not fully open (**Figure 1D-E**). The channel deviates significantly from TRPV2_APO_1_ with an overall Cα RMSD of 1.8 Å (**Figure 1F**), yet bears a marked resemblance to our recent WT rat TRPV2 CBD-bound state 2 (TRPV2_CBD_2_, PDB 6U88)^26^ with an overall Cα RMSD of 0.6 Å (**Figure 1G**). Compared to TRPV2_APO_1_, the S4-S5 linker of TRPV2_2APB_ has shifted towards S5 by 2.2 Å and has rotated 10° vertically and 9° laterally (**Figure 1F**), with corresponding shifts to domains that interact with the S4-S5 linker like the TRP helix, which shifts laterally by 2.3 Å. These are similar to the shifts that TRPV2_CBD_2_ makes, and also like that structure the C-terminus of TRPV2_2APB_ is wrapped around the β-sheet region rather than forming a helix like in TRPV2_APO_1_. The major deviation between TRPV2_CBD_2_ and TRPV2_2APB_ is seen at S5, where in the TRPV2_2APB_ structure, the bottom portion of S5 is bent away from the neighboring monomer’s S4-S5 linker by 21° relative to TRPV2_CBD_2_ (**Figure 1G**), creating a gap that is occupied by density we have attributed to 2-APB (**Figure S5**).

Thus, we have identified a novel and TRPV2-specific binding site for 2-APB, but it is currently not possible to determine if we have captured TRPV2 in a closed pre-open state or an inactivated/desensitized state.

### Doxorubicin permeation through TRPV2 in the presence of 2-APB

To further explore how TRPV2 permeates large organic cations, we used 1 mM 2-APB to activate WT rat TRPV2 in nanodiscs in the presence of 5 μM DOXO and obtained a cryo-EM structure (TRPV2_DOXO_) with an overall resolution of 4.0 Å (**Figure S8-S9, Table S1**). We did not apply symmetry while refining the TRPV2_DOXO_ map due to the expected asymmetric position of DOXO, yet the structure of the TRPV2 channel itself maintained C4 symmetry. A comparison of TRPV2_DOXO_ to TRPV2_2APB_ revealed that the structures are almost identical with an overall Cα RMSD of 0.6 Å. Thus, we have captured 2-APB-bound TRPV2 but with an additional large non-protein density in the middle of the pore just below the selectivity filter (**Figure 2A-B, Figure S10**). This new density occupies the widest part of the pore (**Figure 2C-D**) and it is unique to TRPV2_DOXO_; no density of this prominence is observed in the TRPV2_2APB_ map without symmetry (**Figure S10**). The DOXO molecule can be fitted into this density in two opposite orientations (**Figure S11**) and in both cases it seems to be partially stabilized in the proposed positions through interactions between Tyr634 and an amino group on the tetrahydropyranyl fragment of DOXO (**Figure 2B**). While the selectivity filter of this structure seems not to be sufficiently open to allow for the passage of DOXO (**Figure 2C-D**), data from our previous WT rat TRPV2 apo state 2 structure (TRPV2_APO_2_, PDB 6U86)^26^ and recent electrophysiological studies^57^ show that the selectivity filter is quite dynamic and could easily allow DOXO to enter the pore (**Figure 2D**). The lower gate of the channel remains closed at Met645 (**Figure 2C-D**), which may have been why we were able to trap DOXO inside the pore. Thus, our data shows that the DOXO cation can penetrate the selectivity filter of TRPV2 in the presence of 2-APB and is sufficiently stable inside the pore to allow visualization with single-particle cryo-EM.

**Figure 2.**
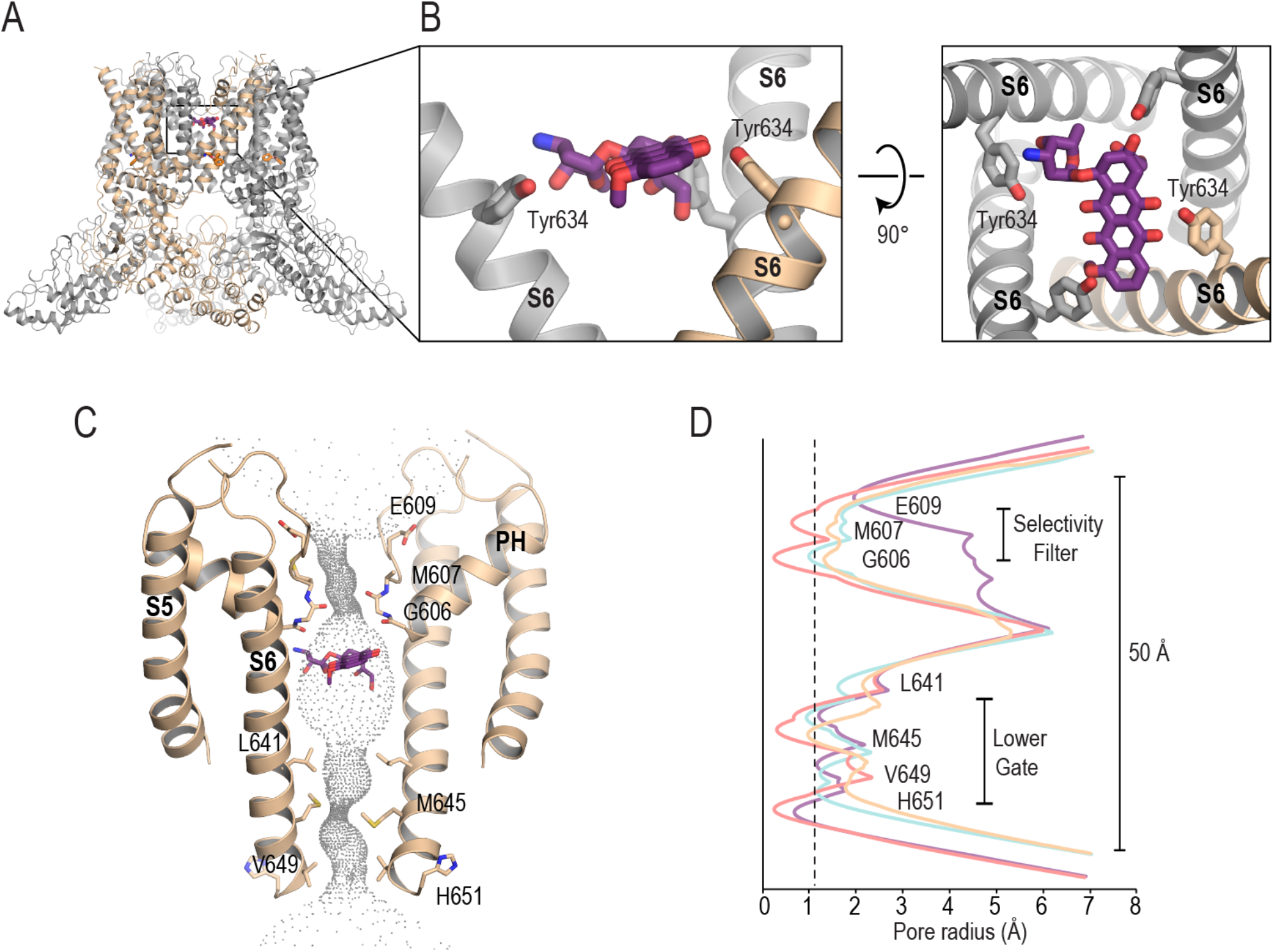
DOXO dwells transiently in the pore. (A) Cartoon representation of the TRPV2_DOXO_ model. (B) Detailed view a possible position for DOXO in the pore; DOXO is depicted as purple sticks. (C) Pore profile of TRPV2_DOXO_ and (E) a graphical representation of the pore profiles of TRPV2_DOXO_ (wheat), TRPV2_2APB_ (pale cyan), TRPV2_APO_1_ (PDB 6U84) (salmon), and TRPV2_APO_2_ (PDB 6U86) (purple).

### TRPV2 pore block by ruthenium red

To uncover the structural basis for TRPV2 pore block by RR, we incubated WT rat TRPV2 reconstituted in nanodiscs with 10 μM RR for 5 minutes before preparing grids for cryo-EM. The resulting dataset yielded a structure of RR-bound TRPV2 (TRPV2_RR_) with C4 symmetry at an overall resolution of 3.1 Å (**Figure S12-13**, **Table S1**). The structure revealed that RR stabilized TRPV2 in a state almost identical to TRPV2_APO_2_ with a Cα RMSD of 0.7 Å^26^. Notably, the TRPV2_APO_2_-like state was the dominant conformation in the RR dataset, while our apo TRPV2 dataset contained predominantly particles in the TRPV2_APO_1_ state^26^. After examination of non-protein densities present in this structure, we have identified a putative RR binding site at the junction between the bottom of the pore helix and the pore loop (**Figure 3A-B, Figure S14**). At this position, the RR molecule is coordinated by the backbone carbonyls of residues of the selectivity filter (Thr604-Gly608) from adjacent monomers, while the sidechains of Phe603 and Tyr634 could stabilize RR with cation-π interactions. **(Figure 3B)**. A carboxyl group of Glu609 is positioned above the RR molecule and could be contributing to its stabilization in the binding pocket (**Figure 3B**). Glu609 equivalents in TRPV1 (Asp646), TRPV3 (Asp641), and TRPV4 (Asp682) impact efficacy of pore blockade by RR^58–60^, suggesting that the mechanism of pore blocking by RR is conserved in TRPV1-4 channels.

**Figure 3.**
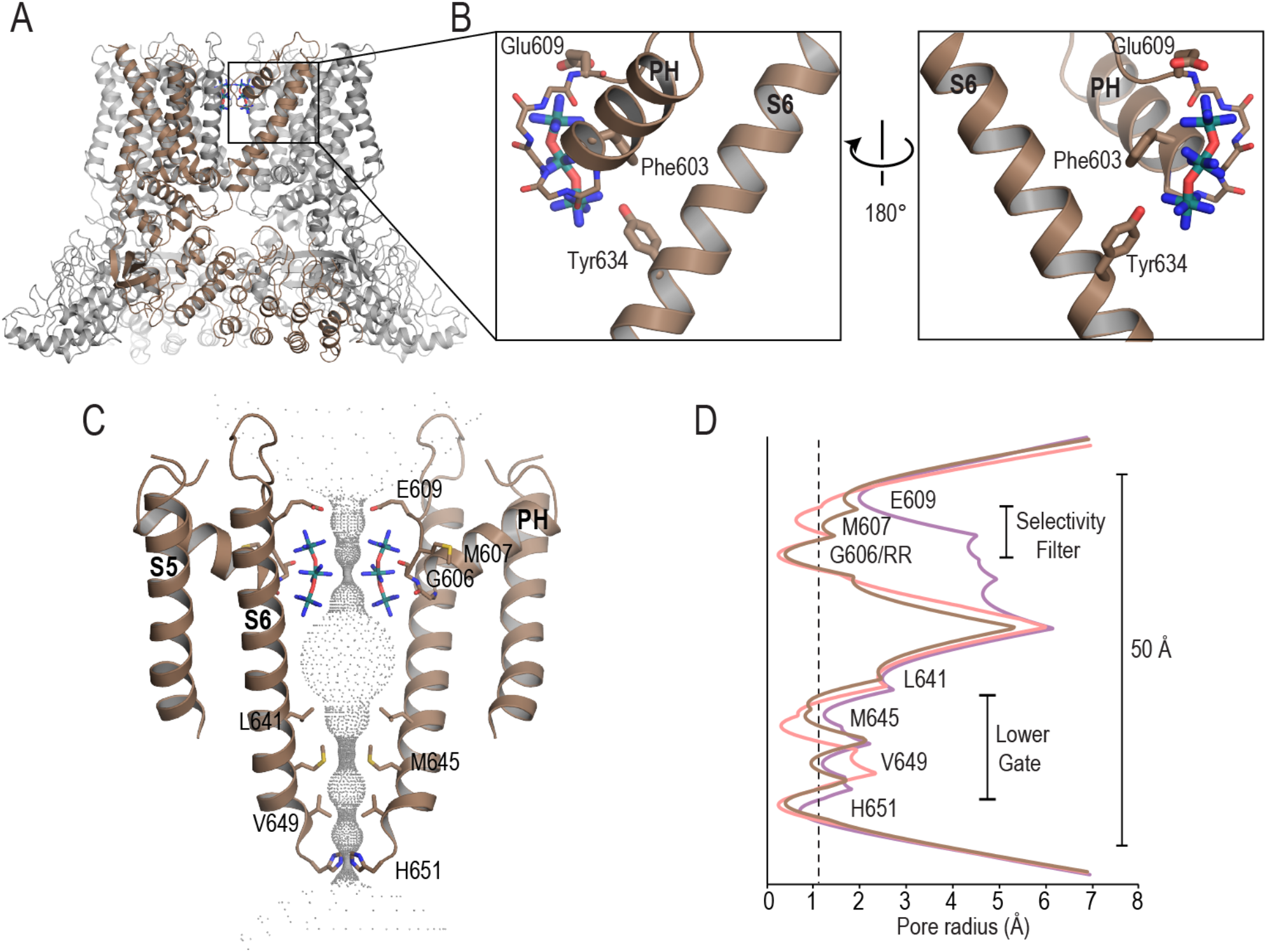
RR binds to the selectivity filter and blocks the pore. (A) Cartoon representation of the TRPV2_RR_ model. (B) Detailed view of the RR binding site; RR is depicted as red sticks. (C) Pore profile of TRPV2_RR_ and (E) a graphical representation of the pore profiles of TRPV2_RR_ (brown), TRPV1_APO_1_ (PDB 6U84) (salmon), and TRPV2_APO_2_ (PDB 6U86) (purple). RR is depicted as red sticks.

The proposed binding site suggests a robust mechanism of pore blocking by RR. Despite the opened selectivity filter, similar to TRPV2_APO_2_ (**Figure 3D**), four molecules of RR completely block the ion permeation pathway at the top of the pore (**Figure 3C**), narrowing the radius of the selectivity filter of the channel to the level of TRPV2_APO_1_ (**Figure 3D**). In addition to steric hindrance, positively charged RR creates an electrostatic barrier for cations entering the pore. Since the selectivity filter undergoes spontaneous opening in the apo protein^26,57^, the binding site could be easily accessible for RR. Critically, the inherent flexibility of the selectivity filter would mean that pore blocking at this site could be completely independent from channel opening^26,57^.

The TRPV2_RR_ structure had significantly higher resolution compared to the original TRPV2_APO_2_, resolved at 4.0 Å^26^, allowing us to improve the model, especially in the area of S4-S5 linker where we were able to more clearly identify density for a structural lipid in the vanilloid pocket. The lipid headgroup is positioned between the S4-S5 linker on one TRPV2 subunit and the bottom of S6 on the neighboring subunit (**Figure 4A, Figure S15**). It is not possible to definitively identify the head group of the lipid from this density, but it seems likely that the lipid could be a phosphatidylethanolamine or phosphatidylcholine based on the negatively charged environment created by Asp536, Glu647, and Asp654 that are exposed in this pocket (**Figure S16A**). The phosphate group of the lipid would be coordinated by the backbone amide groups of Gln530 and Ile529, and potentially the amine from Asn639 (**Figure S16B**). The acyl tails of the lipid extend into the vanilloid binding pocket, similar to the lipids observed previously in TRPV1^29^ and TRPV3^40,41^ (**Figure S17**). The same acyl tail densities are present in our earlier TRPV2_APO_1_ and TRPV2_APO_2_ structures^26^ (**Figure 4B, Figure S15**). However, in those structures we were not able to identify the density for the headgroup, which presumably merged with the S4-S5 linker density at 4 Å resolution. Importantly, TRPV2_CBD_2_^26^ had only a small amount of non-protein density in this pocket and TRPV2_2APB_ has none at all despite being at a similar nominal resolution as TRPV2_APO_1_ (**Figure 4C, Figure S15**). Thus, our data suggests that the transition from an apo state to a 2-APB-bound state requires the expulsion of a lipid from the vanilloid pocket. While 2-APB does not bind to the vanilloid pocket, this is comparable to the TRPV1 activation model where a lipid bound in the vanilloid pocket must be displaced before the channel can open^29^.

**Figure 4.**
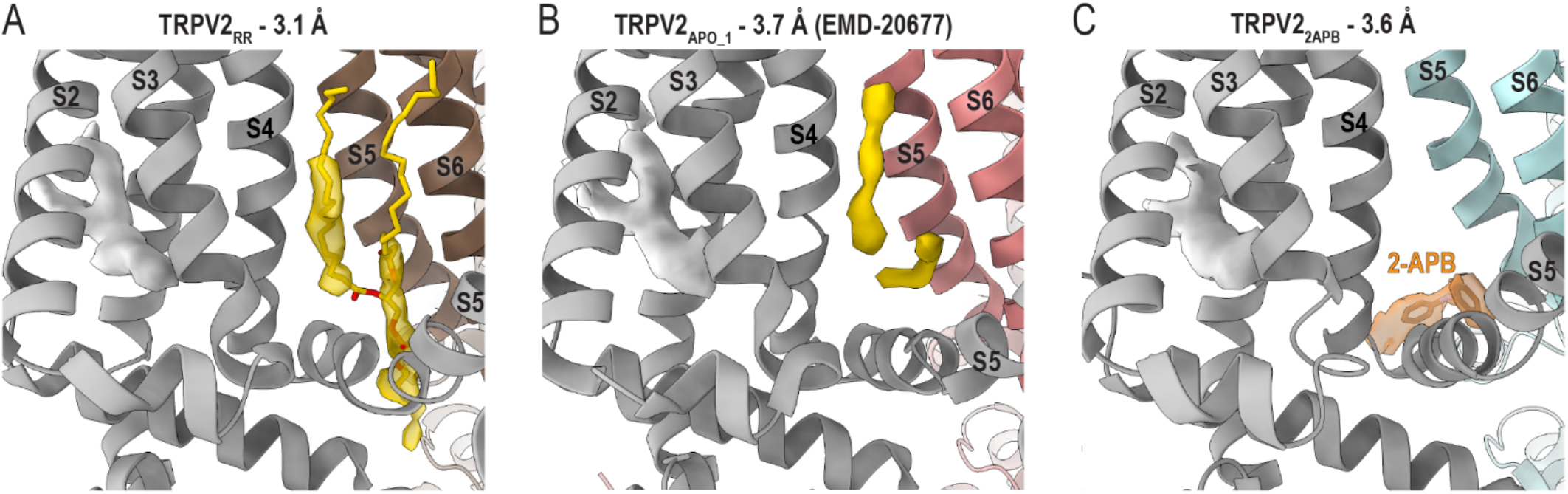
The TRPV2 vanilloid pocket. Cartoon representations of the (A) TRPV2_RR_, (B) TRPV2_APO_1_ (PDB 6U84, EMD-20677), and (C) TRPV2_2APB_ models with cryo-EM map densities for the vanilloid lipid (yellow), the VSLD lipid (white) and 2-APB (orange) depicted as surfaces. The proposed vanilloid lipid is modelled as phosphatidylethanolamine and depicted as yellow sticks; 2-APB is depicted as orange sticks.

## Discussion

Here we have presented WT rat TRPV2 structures in the presence of 2-APB and RR in lipid membranes, which revealed a unique 2-APB binding site for TRPV2 and a conceivably conserved mechanism of TRPV1-4 channel block by RR. We revealed that large organic molecules, like DOXO, can enter the pore of TRPV2 upon its activation by 2-APB, which is consistent with previous observations using cell-based assays for TRPV2 and other TRP channels^13,47,48,52–54^. Additionally, we identified an alternate lipid binding site in the vanilloid pocket of TRPV2 that may play a role in channel modulation by 2-APB.

Neither in this current study using 2-APB as an activator, nor in our previous work with CBD^26^, were we able to capture WT TRPV2 in a fully open state. This could be due to the fast inactivation of 2-APB and the slow activation of CBD (**Figure S2**), or because of cryo-EM experimental conditions. Nevertheless, the TRPV2_2APB_ structure revealed a novel 2-APB binding site, which differs from the 2-APB binding site observed for TRPV3^41–44^. Studies investigating the interaction of 2-APB with TRPV3 pinpointed human His426, located at the pre-S1 helix and tucked against the TRP helix, as a critical residue for activation^41–44^. The 2-APB binding site determined here for TRPV2 also features a key histidine residue, rat His521, which is located at the N-terminal end of the S4-S5 linker, wedged between the S4-S5 linker of one monomer and S5 of an adjacent monomer. His521 is highly conserved in mammalian TRPV2 (**Figure S1B**), but notably, neither it nor His426 in TRPV3 are conserved in other TRPV channels in humans (**Figure S1A**), which suggests that TRPV1 may yield yet another unique 2-APB binding site^44^. It is quite interesting that structurally similar TRPV channels can be activated by the same molecule with completely different binding modes and therefore presumably different mechanisms as well.

To date, TRPV2 structures in the presence of CBD or 2-APB have revealed two entirely unique sites for binding and potential mechanisms for channel activation compared to other TRPV channels. A common feature of these binding sites is that they reside at an interface between channel subunits, which fits with what we know about the inter-subunit interactions necessary for channel opening for TRP channels^1^. Although 2-APB does not activate TRPV2 by binding in the vanilloid pocket, it still appears to displace the vanilloid lipid present in the apo structures, which caused the S4-S5 linker to adopt a continuous helical conformation. This may indicate that the expulsion of the vanilloid lipid is necessary for TRPV2 channel opening, much like in TRPV1^29^. While we are still waiting to see a fully opened TRPV2 structure, it seems likely from this study and earlier works^26,30,48,61^ that the S4-S5 linker is the right target area for development of TRPV2-specific activators.

Upon activation with 2-APB TRPV2 becomes permeable to large organic cations^13,47,48^. Uptake of green-fluorescent dye Yo-Pro^48^ is characterized by slow kinetics, suggesting that either the probability of getting into the pore is low for large organic cations, or that they get stabilized inside the pore and go through slowly, or both. Despite not being able to resolve TRPV2 in a fully open state, our TRPV2_DOXO_ shows that the DOXO cation can penetrate the selectivity filter and make transient contacts with polar groups inside the pore below the selectivity filter. Since DOXO does not function as a channel blocker, contacts inside the pore are not expected to be strong enough to hold DOXO in place and prevent passage through the pore or washing out from the pore. Such transient stabilization is consistent with reversable pore blocking by DOXO that has been observed previously^62^. The lower gate of TRPV2_DOXO_ remains closed like the lower gate of TRPV2_2APB_ reflecting the same functional state of the channel.

While large organic cations can pass through the pore of several TRP channels, the highly positively charged RR is a known TRP channel pore blocker. In this study we were able to identify a new binding site for RR which provides important insights into the channel function. Four RR molecules are inserted in the selectivity filter of TRPV2 while the filter is in a wider conformation, as previously observed in TRPV2_APO_2_^26^. This binding mode of RR constricts the selectivity filter sufficiently to prevent ion permeation, although the selectivity filter is not thought to act as a gate^57^. The wider conformation of the selectivity filter in this state exposes a large number of backbone carbonyls which coordinate with RR. Moreover, the acidic residue Glu609 may also coordinate with RR and an acidic residue is conserved in this position among human TRPV1-TRPV4 (**Figure S1A**), suggesting that the mechanism of pore block by RR is conserved in TRPV1-4^58–60^.

Together, our results allow us to propose models for TRPV2 gating. The TRPV2 selectivity filter is not a gate^26,57^, but instead has the conformational flexibility to allow for the entry of large organic cations like DOXO and the binding of pore blockers like RR (**Figure 5**). The binding of 2-APB at the interface between the S4-S5 linker of one monomer and S5 of an adjacent monomer may lead to channel opening by displacing the lipid bound in the vanilloid pocket in the apo/blocked states of the channel (**Figure 5**). While the fully open state of TRPV2 has yet to be seen, our results reveal new insights into TRPV2 function and will allow for TRPV2-specific drug design.

**Figure 5.**
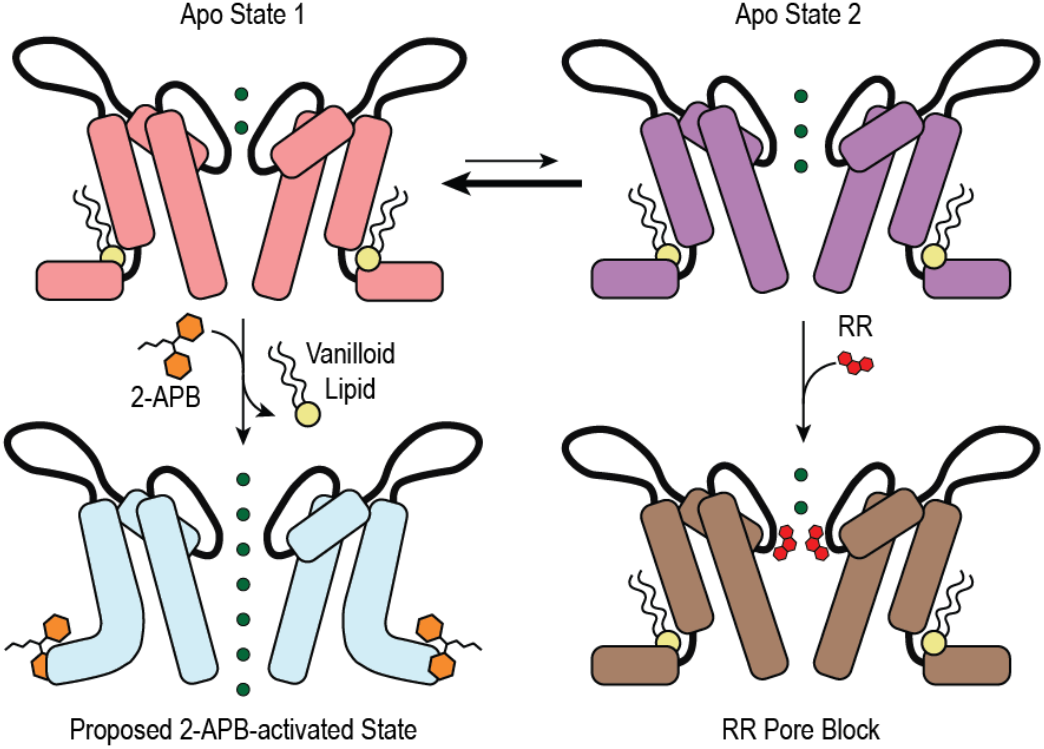
Proposed model of TRPV2 gating. Schematic representations of proposed models for ligand activation and inhibition of TRPV2. In lipid membranes, apo TRPV2 has a lipid in the vanilloid pocket and can transition between Apo State 1 and Apo State 2. The addition of 2-APB expels the lipid from the vanilloid pocket, allowing the channel to open. RR can enter the selectivity filter in Apo State 2 and bind there to prevent ion passage.

## Methods

### Protein sequence alignment of the TRPV channels

TRPV sequence alignments were performed using Clustal Omega^63^. Graphics of sequence alignments were generated using Aline^64^.

### Protein expression and purification

Full-length rat TRPV2 was expressed and isolated as previously published^26^. Briefly, rat TRPV2 cloned into a YepM vector and tagged with 1D4 was expressed in Saccharomyces cerevisiae. The yeast expressing TRPV2 were lysed in 25 mM Tris, pH 8.0, 300 mM Sucrose, and 5 mM EDTA using a M110Y Microfluidizer (Microfluidics), and the membranes isolated by ultracentriguation at 100,000 x g. These TRPV2-enriched membranes were solubilized in 20 mM Hepes, pH 8.0, 150 mM NaCl, 5% glycerol, 0.087% LMNG, 2 mM TCEP, and 1 mM PMSF for 1 hour. This mixture was clarified by centrifugation at 100,000 x g and the insoluble fraction discarded. The solubilized TRPV2 was bound to 1D4 antibody coupled CnBr-activated Sepharose beads, followed by washes with Wash Buffer (20 mM Hepes, pH 8.0, 150 mM NaCl, 2 mM TCEP) supplemented with 0.006% (w/v) DMNG. The protein was eluted with Wash Buffer supplemented with 0.006% (w/v) DMNG and 3 mg/mL 1D4 peptide. The purified TRPV2 was concentrated to a volume under 1 mL and reconstituted into nanodiscs in a 1:1:200 ratio of TRPV2:MSP2N2:soy polar lipids (Avanti). The soy polar lipids were rapidly dried under a nitrogen flow and further dried under vacuum before being resuspended in Wash Buffer containing DMNG in a 1:2.5 ratio (soy polar lipids:DMNG). The assembled nanodisc reconstitution mixture was incubated at 4°C for 30 minutes before adding Bio-Beads to the mixture. After 1 hour, the reconstitution mixture was transferred to a new tube with fresh Bio-Beads and incubated overnight at 4°C. The nanodisc embedded TRPV2 was purified from empty nanodiscs using a Superose 6 column (GE) equilibrated in Wash Buffer for size-exclusion chromatography. The eluted TRPV2 was concentrated to 2 mg/mL to use in vitrification.

MSP2N2 was expressed and purified as previously described^37^. Briefly, MSP2N2 inserted into a pET28a vector (Addgene) was expressed in BL21 (DE3) cells. After harvest, the cells were resuspended in a buffer containing 20 mM Tris-HCL, pH 7.4, 1 mM PMSF, and a complete EDTA-free protease inhibitor cocktail tablet (Roche) and lysed using a M110Y Microfluidizer (Microfluidics). The lysate supernatant was bound to Ni-NTA resin, which was then washed with Wash Buffer (20 mM Tris-HCl, pH 7.4, 100 mM NaCl), then by Wash Buffer supplemented with 1% Triton X-100, then by Wash Buffer supplemented with 50 mM sodium cholate, then by Wash Buffer supplemented with 20 mM imidazole. MSP2N2 was then eluted from the Ni-NTA resin with Wash Buffer supplemented with 300 mM imidazole and exchanged into 50 mM Tris-HCl, pH 7.5, 100 mM NaCl, 5 mM EDTA before being concentrated to ~10 mg/mL.

### Cryo-EM sample preparation

For 2-APB-bound TRPV2, 1 mM 2-APB was incubated with TRPV2 for 30 minutes on ice prior to grid preparation. For the 2-APB + DOXO sample, 1 mM 2-APB and 5 μM DOXO were incubated with TRPV2 for 5 minutes prior to grid preparation. For the RR-bound sample, 10 μM RR was incubated with TRPV2 for 5 minutes prior to grid preparation. Immediately before blotting, 3 mM fluorinated Fos-choline 8 (Anatrace, USA) was added to improve particle distribution in vitreous ice. 3 μL of sample was applied to a freshly glow discharged 200 mesh Quantifoil 1.2/1.3 grid and then blotted for 5-8 seconds at 4°C and 100% humidity before vitrification in liquid ethane.

### Cryo-EM data collection

All cryo-EM images were collected on the same 300kV Titan Krios microscope equipped with a Gatan K3 direct detector camera in super resolution mode. All images were collected as 40 frames movies with a total dose of 45 e^-^/A^2^ and a super-resolution pixel size of 0.53 Å/pix. Defocus values ranged from −0.8 to −3.0 μm.

### Cryo-EM data processing

All data processing was done using Relion 3.1. For all three datasets, the movies were motion corrected using the Relion implementation of MotionCor2, binning the pixels to 1.06 Å/pix. The defocus values for the resulting micrographs were estimated using CTFFIND4^65^.

For TRPV2_2APB_, an initial round of manual picking and 2D classification yielded templates for auto-picking covering multiple particle orientations. Template based auto-picking yielded an initial set of 1,350,612 particles. These particles were binned by 4 during extraction, to a pixel size of 4.24 Å/pix, and subjected to 2D classification. The best 1,076,973 particles from 2D classification symmetry and subjected to CTF aberration and anisotropy correction and defocus refinement before 3D classification into 4 classes without angular assignment. Two minor classes comprising 11% of the particles had low resolution, the remaining 89% were distributed between two classes with good TRPV features and continuous helices in the transmembrane domain. However, only one class with 75,691 particles, or 22% of the input, had a big non-protein density in the pore. Those particles were refined without imposing a symmetry, classified to improve resolution and 60,623 best particles were subjected to Bayesian polishing. After the final round of 3D classification, we got the final subset of 58,658 particles. The final pixel size was adjusted to 1.07 Å/pix based on comparison with a TRPV2 ARD crystal structure (PDB 2ETA)^66^ and the map was sharpened in Relion to 4.0 Å.

For TRPV2_RR_, templates for auto-picking were obtained through 2D classification of particles picked by Laplacian-of-Gaussian auto-picking on a subset of micrographs. Template based autopicking from the whole dataset yielded an initial set of 1,498,726 particles. These particles were binned by 4 during extraction, to a pixel size of 4.24 Å/pix, and subjected to two rounds of 2D classification. The best 658,171 particles from 2D classification were unbinned to 2.12 Å/pix, recentered, and classified in 3D with initial angular assignment using the TRPV2_APO_1_ map (EMD-20677) lowpass filtered to 60 Å as an initial reference. 84% of those particles sorted into 5 classes with real TRP channel features, however two of those classes (31% of particles) had strong C2 symmetry, while the remaining three classes had C4 symmetry and were different from each other only in the resolution. For further processing we picked the most high-resolution class containing 150,212 particles, or 23% of the input. Those particles were unbinned to 1.06 Å/pix, refined without symmetry, and classified into three classes without angular assignment, allowing us to identify a 67,031 best particles in a high-resolution class in Apo2-like conformation with additional non-protein density at the selectivity filter. We imposed C4 symmetry on the final 67,031 set of particles, subjected it to CTF aberration and anisotropy refinement followed by two rounds of defocus refinement and Bayesian polishing. The final pixel size was adjusted to 1.07 Å/pix based on comparison with a TRPV2 ARD crystal structure (PDB 2ETA)^66^ and the map was sharpened with Phenix Autosharpen^67^ using a resolution cutoff of 3.1 Å.

### Model Building

For TRPV2_2APB_, the model of TRPV2_CBD_2_ (PDB 6U88) was used as a starting model and docked into the TRPV2_2APB_ map in Coot. For TRPV_2DOXO_, the model of TRPV2_2APB_ was used as a starting model and docked into the TRPV2_DOXO_ map in Coot. For TRPV2_RR_, the model of TRPV2_Apo_2_ (PDB 6U86) was used as a starting model and docked into the TRPV2_RR_ map in Coot. These models were then iteratively manually adjusted to the map and refined using phenix.real_space_refine from the PHENIX software package.

The ligand restraint files for 2-APB, DOXO, and RR were generated using the eLBOW tool from the PHENIX software package^68^. Images of the models and maps for figures were generated using Pymol (Schrödinger, USA), Chimera^69^, and ChimeraX^70^.

### Inside-out single channel recordings

WT rat TRPV2 cDNA was transiently transfected into HEK293T cells and seeded on 60 mm petri dishes to high confluence for performing single channel measurements in the inside-out configuration, much as previously described^71^. Glass borosilicate capillaries (#30-0076, MultiChannel Systems MCS, Reutlingen, Germany) were pulled with a resistance of 20-50 MΩ and coated with Sigma Cote ® (Sigma-Aldrich, Munich, Germany) to reduce their capacitance. Currents were measured with CBD and 2-APB applied by a gravity-driven perfusion system and recorded at +60 mV at room temperature. The external solution contained: 140 mM KCl, 5 mM NaCl, 2mM MgCl_2_, 5 mM EGTA, and 10 mM HEPES, pH 7.4 (adjusted with KOH). The pipette solution contained: 140 mM NaCl, 5 mM KCl, 2 mM MgCl_2_, 5 mM EGTA, 10 mM glucose, and 10 mM HEPES, pH 7.4 (adjusted with NaOH). The liquid junction potential (−3.8 mV) was calculated and subtracted *a priori*. Current traces were low-pass filtered using an analogue 1 kHz eight-pole Bessel filter and subsequently digitally filtered at 300 Hz. Sampling rate was 10 kHz. Data acquisition and off-line analysis were performed by a combination of Patchmaster/Fitmaster software (HEKA Elektronik), Clampex/Clampfit 10.7 software (Axon, Molecular Devices, Sunnyvale CA, U.S.A.) and Origin 2020 (Origin Lab, Northampton, MA, U.S.A.)

### Whole-Cell Voltage Clamp

Nanofectin (PAA, Pasching, Austria) was used to transfect HEK293T cells with WT rat TRPV2, Arg539Ala rat TRPV2, His521Ala rat TRPV2 or Thr522Ala rat TRPV2. Mutants were generated according to the instructions of the manufacturer by site directed mutagenesis with the quick-change lightning site-directed mutagenesis kit (Aglient, Waldbronn, Germany). All mutants were sequenced to confirm intended amino acid exchange and to exclude further channel mutations. Under standard cell culture conditions (5% CO_2_ at 37°C) cells were cultured at 37°C with 5% CO_2_ in Dulbecco’s modified Eagle medium nutrient mixture F12 (DMEM/F12 Gibco/Invitrogen, Darmstadt, Germany) and supplemented with 10% fetal bovine serum, (Biochrom, Berlin, Germany).

Whole-cell voltage clamp was performed with an EPC10 USB HEKA amplifier (HEKA Electronik, Lamprecht, Germany), much as previously described^71^. Signals were low passed at 1 kHz and sampled at 2 to 10 kHz. Patch pipettes were pulled from borosilicate glass tubes (TW150F-3; World Precision Instruments, Berlin, Germany) to give a resistance of about 3 MΩ. Cells were held at −60 mV and all recordings were performed at room temperature. The external solution contained: 140 mM NaCl, 5 mM KCl, 2 mM MgCl_2_, 5 mM EGTA, 10 mM glucose, and 10 mM HEPES, pH 7.4 (adjusted with NaOH). Calcium was omitted in order to avoid desensitization. The standard pipette solution contained: 140 mM KCl, 2 mM MgCl_2_, 5 mM EGTA and 10 mM HEPES, pH 7.4 (adjusted with KOH). A gravity–driven glass multibarrel perfusion system was used to bath apply solutions. Data acquisition and off-line analyses required Patchmaster/Fitmaster software (HEKA Electronik, Lamprecht, Germany) and Orgin 8.5.1 (Origin Lab, Northampton, MA, U.S.A.).

### Human TRPV2 mutagenesis

The pEGFP-N2 vector containing human TRPV2 fused to FLAG tag and red fluorescent protein was kindly provided by Prof. Kojima (Gunma University, Japan). The gene was further amplified by PCR using the primers 5’-AAAGGGAAGGGAATTATGACCTCACCCTCCAGC-3’ and 5’-CTCGAGCGGCCGCTCTTAGGCGCCGGTGGAGTG-3’, and added to the pcDNA3.1(+) via In-Fusion strategy (Takara Bio, USA), resulting in a plasmid coding for a C-terminal flag-RFP hTRPV2 fusion protein. The products were transformed into *E. coli* NZY5α (Nzytech, Portugal), and successful transformants were isolated on LB agar medium containing 10 μg/ml ampicillin. To produce mutant variants from human TRPV2, the hTRPV2-flag-RFP pcDNA3.1(+) plasmid was used as a template. The His519Ala and Arg537Ala single-point mutations were introduced using the Phusion Site-Directed Mutagenesis Kit (Thermo Fisher Scientific, USA). The mutagenic primers were Arg537Ala forward 5’-CGGGACCTGCTGGCCTTC CTTCTG-3’, Arg537Ala reverse 5’-GGTCCCGCAGGATGACCTTCTGG-3’; His519Ala forward 5’-GTG GCTTCCAGGCAACAGGCATCTA-3’; and His519Ala reverse 5’-AAGCCACGTGTATAGTAAAG CAGGTT-3’.The amplification constructs were transformed into *E. coli* NZY5α. Correct isolates were identified by the expected PCR amplicons from the plasmid constructs and by sequencing performed at the Eurofins Genomics Facility.

### Cell culture and transfection

Human Embryonic Kidney 293T cells (HEK293T, ATCC catalogue # CRL-3216) were cultured to 30% confluency in Dulbecco’s Modified Eagle’s Medium, containing 10% (v/v) fetal bovine serum (FBS, Gibco, USA), 1% (v/v) GlutaMAX (Gibco, USA), 1% (v/v) non-essential amino acids (Gibco, USA), and 1% (v/v) penicillinstreptomycin (Gibco, USA) in 10% CO_2_ at 37°C. For transient transfection, cells were seeded in 8 well μ-plates (Ibidi, Germany) overnight and transfected with WT human TRPV2-flag-RFP pcDNA3.1(+), hTRPV2 mutant variants, or RFP pcDNA3.1(+) empty vector using Fugene HD Reagent (Promega, USA). Calcium imaging was performed after 24-hour incubation in 10% CO_2_ at 37°C.

### Calcium influx imaging

Intracellular calcium measurements were performed with Fura-2 AM calcium indicator (Life Technologies, USA) according to previously published protocol ^72^ with minor modifications. Briefly, one hour before the measurements, cells were loaded with 5 μM Fura-2 AM for 45 min. Then, cell medium was replaced by Tyrode’s solution (119 mM NaCl, 5 mM KCl, 2 mM CaCl2, 2 mM MgCl_2_, 6 g/L glucose, 25 mM HEPES pH 7.4), and cells were allowed to readjust to the new medium for 15 min. Fura-2 AM emissions in response to excitation at 340 nm and 380 nm were recorded for 40 seconds before TRPV2 activation with 1mM 2-APB for 260 seconds. Samples were imaged at room temperature using an inverted epifluorescence microscope Axiovert 135TV (Zeiss, Germany) equipped with a high-speed illumination system Lambda DG4 (Shutter Instrument, USA). Data were recorded by a CCD camera. Images were collected and analyzed using MetaFluor Fluorescence Ratio Imaging Software (Molecular Devices, USA). For statistical analysis, 40 most responsive cells per image were selected manually, based on increase in [Ca^2+^] concentration. Experiments were performed in triplicate.

## Acknowledgements

We acknowledge the use of instruments at the Electron Microscopy Resource Lab and at the Beckman Center for Cryo Electron Microscopy at the University of Pennsylvania Perelman School of Medicine. We also thank Darrah Johnson-McDaniel and Edwin Fluck for assistance with Krios microscope operation at the University of Pennsylvania. We thank Sabine Baxter for assistance with hybridoma and cell culture at the University of Pennsylvania Perelman School of Medicine Cell Center Services Facility. We thank Marta Marques and Sandra Vaz for help in analysis of calcium imaging experiments at IMM. We thank Christine Herzog and Kerstin Reher who created TRPV2 mutant constructs at Hannover Medical School. This work was supported by grants from the National Institute of Health (R01GM103899 and R01GM129357 to V.Y.M.-B.), from the Royal Society (URF\R\180019 to G.J.L.B.), from FCT Portugal (FCT Investigator, IF/00624/2015 to G.J.L.B., SFRH/BPD/143583/2019 to B.B.S.).

## Data Availability

The atomic coordinates and cryo-EM density maps of the structures presented in this paper are deposited in the Protein Data Bank and Electron Microscopy Data Bank under the accession codes PDB 7JO4 and EMD-22405 (TRPV2_2APB_), PDB 7JO5 and EMD-22406 (TRPV_2DOXO_), and PDB 7JO6 and EMD-22407 (TRPV2_RR_).

## Contributions

A.D.P., R.A.P. and V.Y.M.-B. conceived and designed all experiments in the manuscript. A.D.P. R.A.P. and P.N.G. purified protein and prepared cryo-EM samples. A.D.P. and R.A.P. processed cryo-EM data, built structural models, analyzed structural data, and prepared figures. J.d.l.R. and F.M.H. performed and analyzed electrophysiology experiments. B.B.S. created TRPV2 mutant constructs and performed and analyzed calcium flux experiments. G.J.L.B. supervised the calcium flux experiments. A.L. supervised the electrophysiology experiments. V.Y.M.-B. supervised all experiments in the manuscript. A.D.P., R.A.P. and V.Y.M.-B. drafted and revised the manuscript. All authors reviewed and approved the final version of the manuscript.

---

**Table S1.** Cryo-EM data collection and model statistics

|  | TRPV2<sub>2APB</sub><br>(EMD-22405,<br>PDB 7JO4) | TRPV2<sub>DOXO</sub><br>(EMD-22406,<br>PDB 7JO5) | TRPV2<sub>RR</sub><br>(EMD-22407,<br>PDB 7JO6) |
| --- | --- | --- | --- |
| **Data collection and processing** |  |  |  |
| Magnification | 81,000x | 81,000x | 81,000x |
| Voltage (kV) | 300 | 300 | 300 |
| Defocus range (μm) | -0.8 to -3.0 | -0.8 to -3.0 | -0.8 to -3.0 |
| Pixel size (Å) | 1.07 | 1.07 | 1.07 |
| Total extracted particles (no.) | 1,350,612 | 4,815,799 | 1,498,726 |
| Refined particles (no.) | 1,076,973 | 2,256,466 | 658,171 |
| Final particles (no.) | 39,166 | 58,658 | 67,031 |
| Symmetry | C4 | C1 | C4 |
| Map sharpening *B* factor (Å<sup>2</sup>) | -76 | -129 | -52 |
| Map resolution (Å) | 3.6 | 4.0 | 3.1 |
| FSC threshold | 0.143 | 0.143 | 0.143 |
| **Model Refinement** |  |  |  |
| Model resolution cut-off (Å) | 3.6 | 4.0 | 3.1 |
| Model composition |  |  |  |
| Nonhydrogen atoms | 19,428 | 19,228 | 20,244 |
| Protein residues | 2,460 | 2464 | 2,496 |
| Ligands | FZ4: 4 | 0 | R2R: 4 |
| R.M.S. deviations |  |  |  |
| Bond lengths (Å) | 0.006 | 0.007 | 0.008 |
| Bond angles (°) | 0.850 | 1.096 | 0.900 |
| Validation |  |  |  |
| MolProbity score | 1.42 | 1.74 | 1.25 |
| Clashscore | 4.78 | 6.95 | 3.16 |
| Rotamers outliers (%) | 0.20 | 0.05 | 0.18 |
| CαBLAM outliers (%) | 1.50 | 2.75 | 0.82 |
| Ramachandran plot |  |  |  |
| Favored (%) | 97.03 | 94.78 | 97.24 |
| Allowed (%) | 2.97 | 5.22 | 2.76 |
| Disallowed (%) | 0.00 | 0.00 | 0.00 |

**Figure S1.**
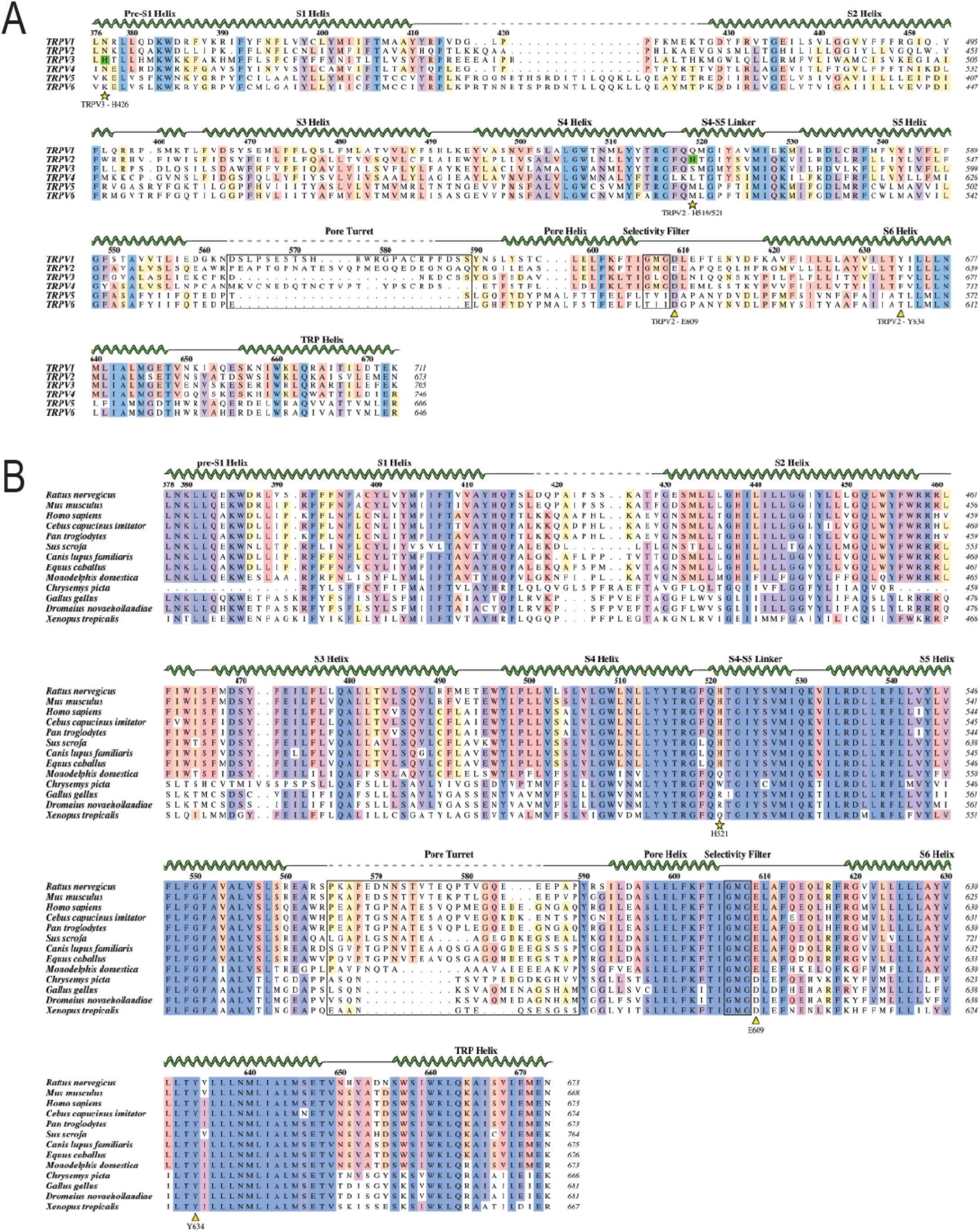
Sequence alignments of the TRPV2 transmembrane domain. Sequence alignments of (A) the human TRPV family and (B) a selection of TRPV2 orthologs from the Pre-S1 helix to the TRP helix. The human TRPV family alignment is numbered based on human TRPV2, the TRPV2 ortholog alignment is numbered based on rat TRPV2, residue numbers for each sequence are marked at the end of each line. Residues with conserved identity are highlighted with a gradient from pale yellow (50%) to blue (100%). The key histidine residues involved in 2-APB interaction with TRPV2 and TRPV3 are marked with a green highlight in the human TRPV family alignment.

**Figure S2.**
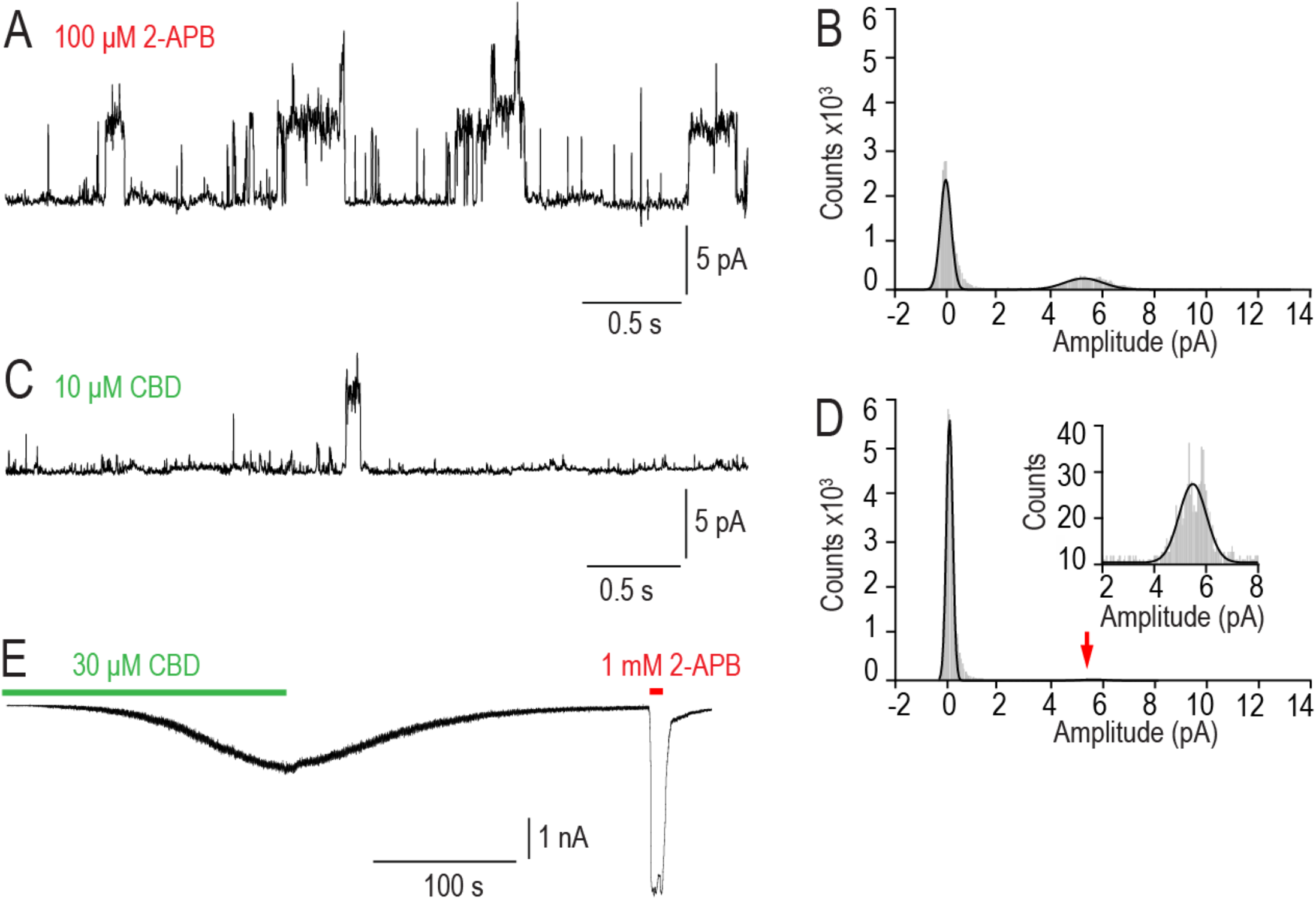
2-APB and CBD responses from HEK293T cells expressing rat TRPV2. Representative inside-out single channel recordings of WT rat TRPV2 during application of 100 μM 2-APB (A) or 10 μM CBD (C). The shown traces are a 4 second section after >2 min application of CBD, which was subsequently washed out, followed a 4 second section after >30 s application of 2-APB. Amplitude histograms for both 2-APB and CBD single channel traces (B and D). The amplitude histograms show 7% absolute open probability for CBD and 21% absolute open probability for 2-APB. The holding potential was set at +60 mV. Recordings were performed in gap-free mode. (E) Whole-cell voltage clamp recording on WT rat TRPV2 at −60 mV displaying inward currents evoked by 30 μM CBD followed by 1 μM 2-APB.

**Figure S3.**
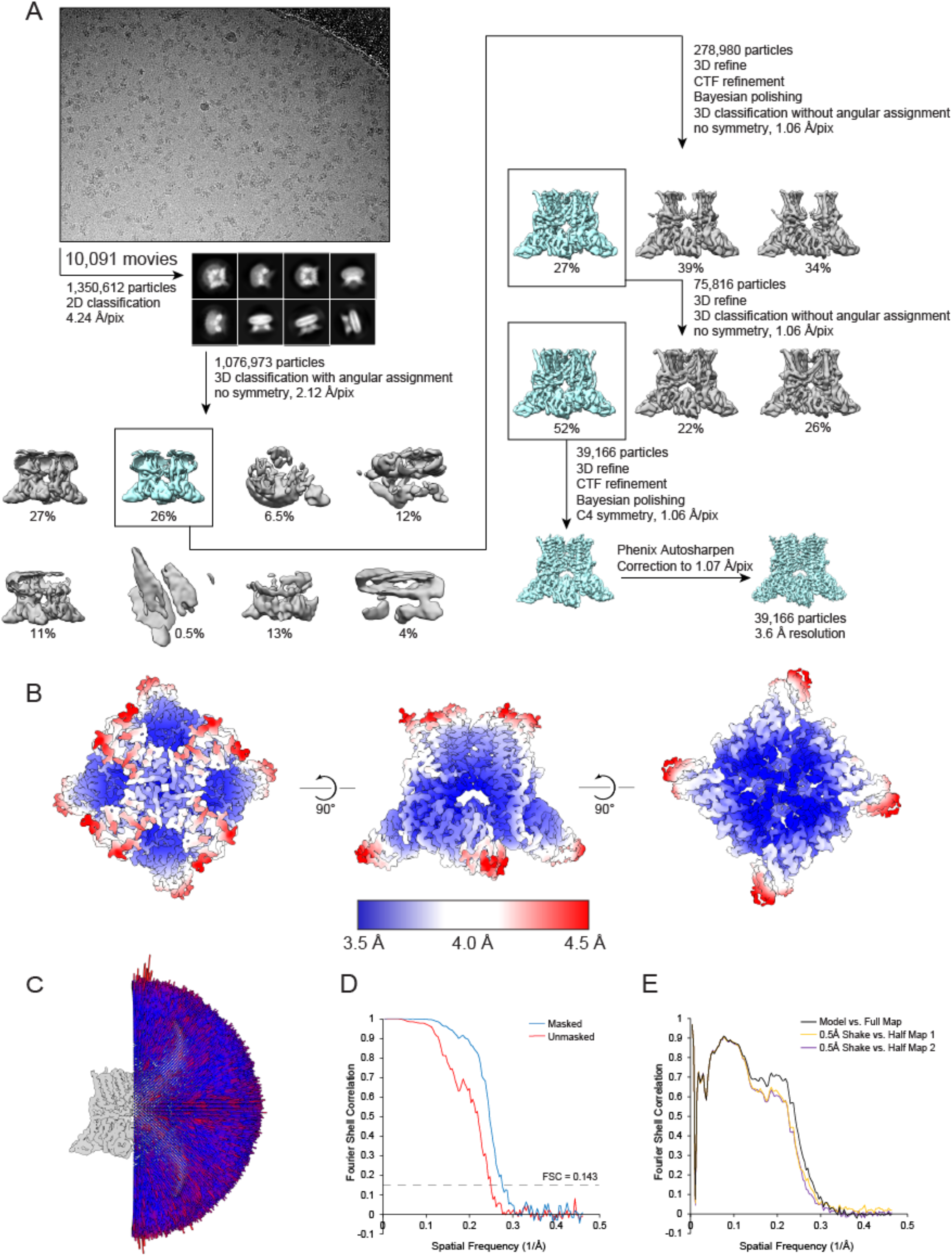
Overview of cryo-EM data from TRPV2_2APB_. (A) Diagram of the data processing path from Relion. (B) Local resolution of the TRPV2_2APB_ map. (C) Angular distribution of particles. FSC curves of the final map (D) and model validation (E).

**Figure S4.**
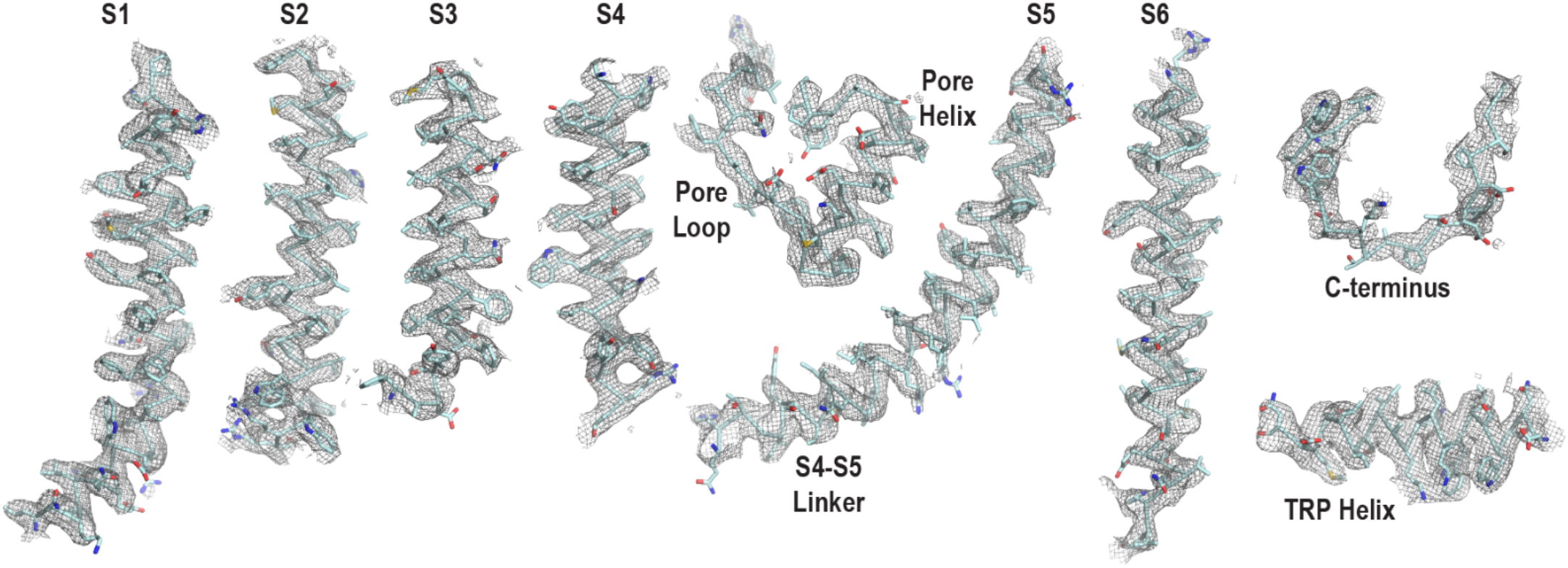
Representative densities of the TRPV2_2APB_ cryo-EM map. Model is depicted as a ribbon with sidechains as sticks; densities are contoured at σ = 4.5.

**Figure S5.**
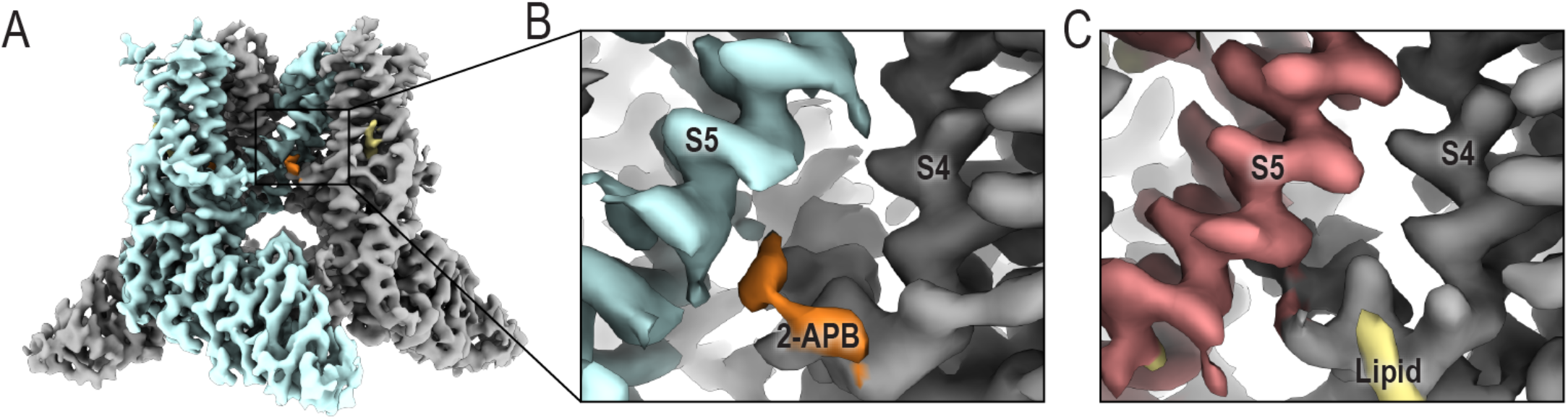
Cryo-EM density at for 2-APB at the S4-S5 linker. (A) Surface representation of the TRPV2_2APB_ cryo-EM map. Detailed view of (B) the 2-APB binding site and (C) the same site from TRPV2_APO_1_ (EMD-20677). One monomer of TRPV2 is colored grey, the adjacent monomer is colored pale cyan (TRPV2_2APB_) or salmon (TRPV2_APO_1_); density attributed to 2-APB is colored orange, density attributed to lipid is colored khaki.

**Figure S6.**
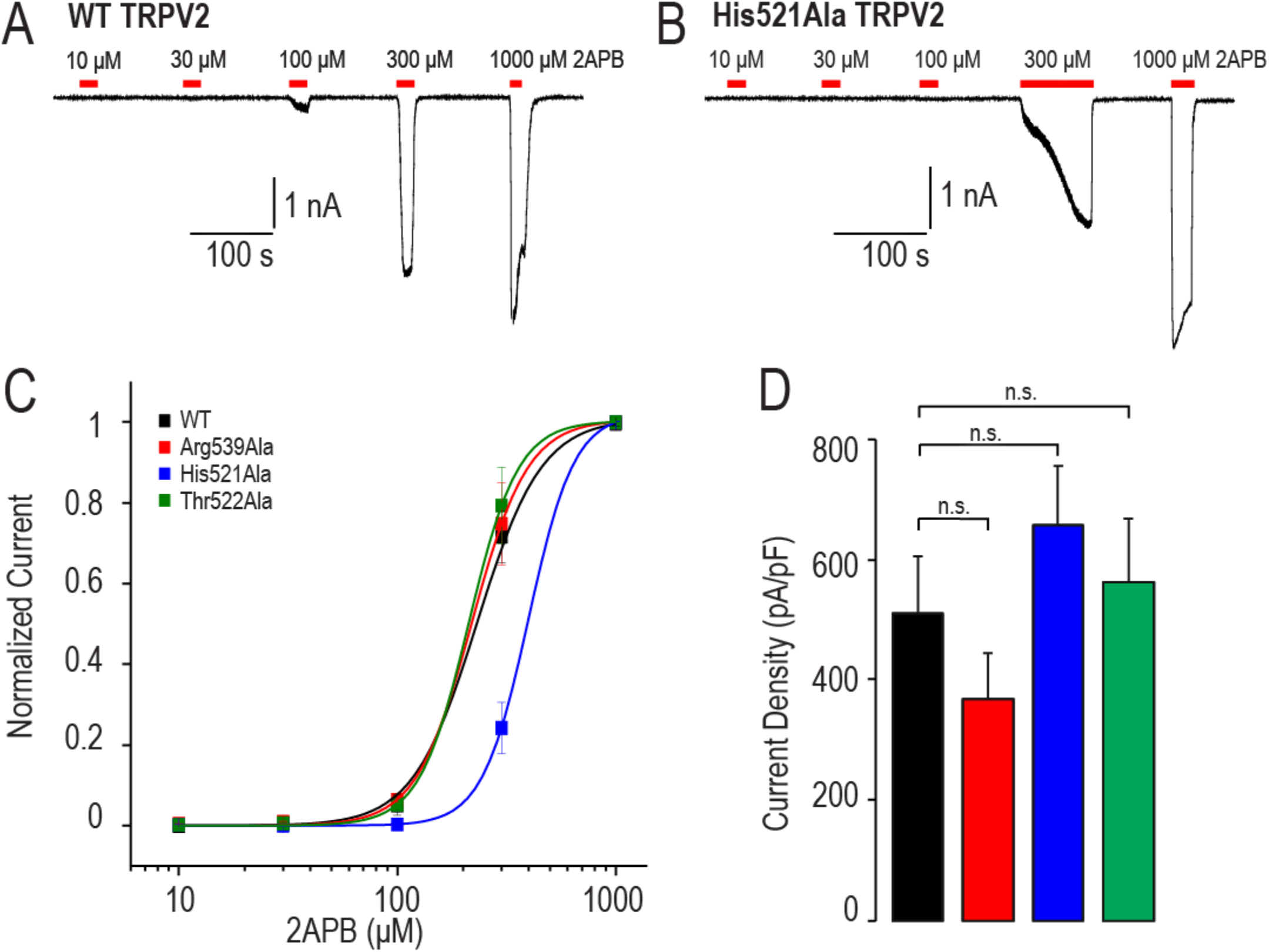
Whole-cell patch clamp recordings of HEK293T cells expressing rat TRPV2. Representative whole-cell patch clamp current traces displaying a concentration-dependent activation of WT rat TRPV2 (A) and His521Ala rat TRPV2 (B) by 2-APB. Cells were held at −60 mV and increasing concentrations of 2-APB were applied. (C) Dose-response curves for 2-APB-evoked activation of WT rat TRPV2 (black, EC_50_=232±20 μM n=10), Arg539Ala rat TRPV2 (red, EC_50_=219±27 μM, n=6), His521Ala rat TRPV2 (blue, EC_50_=398±51 μM, n=8) and Thr522Ala rat TRPV2 (green, EC_50_=212±28 μM, n=5). Current amplitudes were determined for each concentration and normalized to the maximal amplitude obtained with 1000 μM 2-APB. The solid lines represent fits calculated with the Hill equation. (D) Bar columns displaying the average current densities of inward current induced by 1000 μM 2-APB on WT rat TRPV2 (black), Arg539Ala rat TRPV2, His521Ala rat TRPV2 (blue) and Thr522Ala rat TRPV2 (green). The current density (pA/pF) was calculated by dividing the absolute current amplitude (pA) with the cell capacitance (pF). Data are shown as mean ± S.E.M. n.s. denotes not significant.

**Figure S7.**
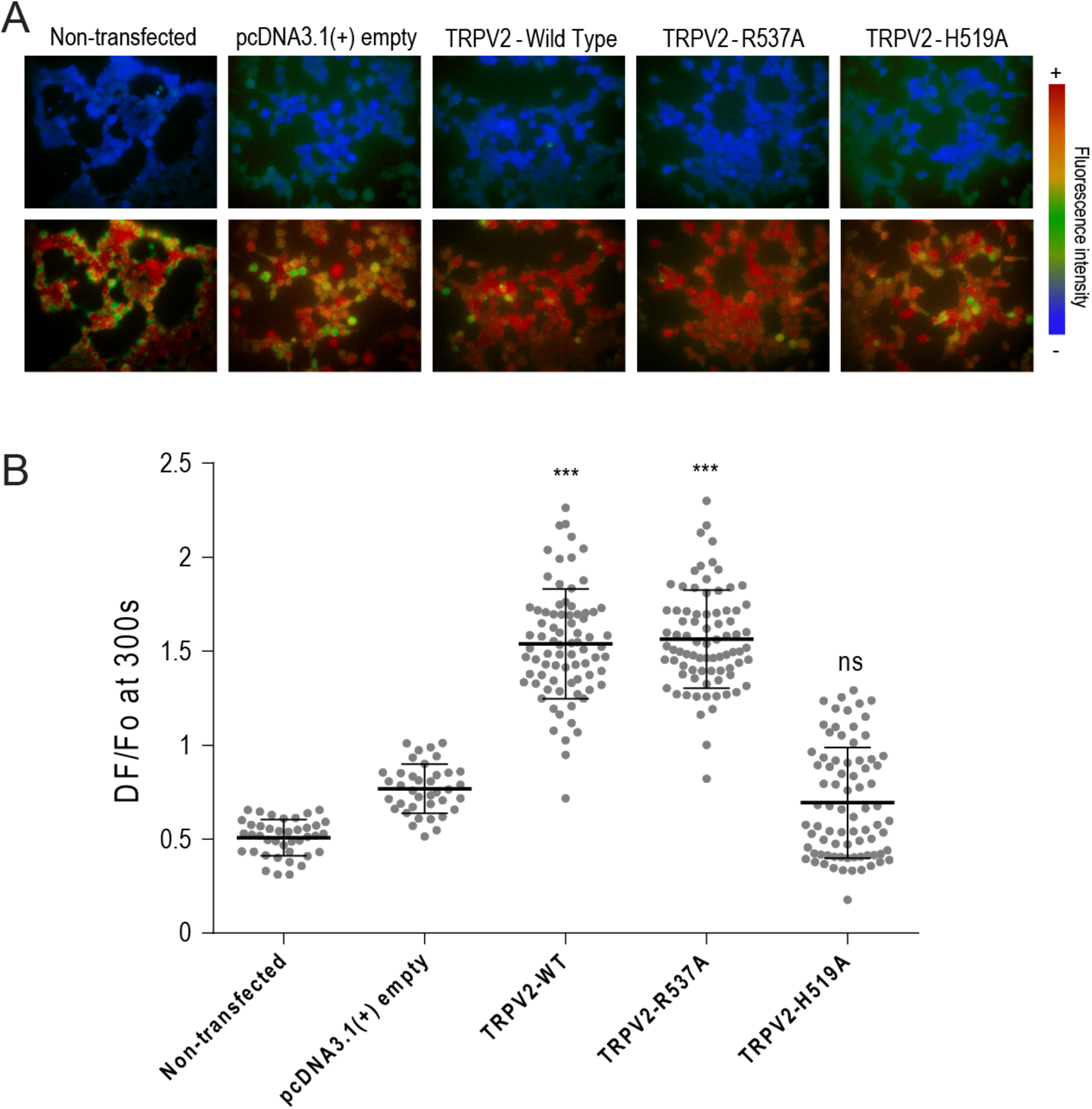
Calcium imaging of HEK293T expressing human TRPV2. Calcium flux imaging of HEK293T cells transiently transfected with WT human TRPV2, Arg537Ala human TRPV2, or His519Ala human TRPV2. Controls of non-transfected cells and cells transfected with pcDNA3.1(+) empty vector are also represented. (A) Representative images taken before (t=40 seconds, top) and after (t=300 seconds, bottom) stimulation with 2-APB. Color palette indicates calcium level (blue – low calcium level; green – intermediate; yellow – medium high; red – high). (B) Quantification of intracellular calcium influx evoked by 2-APB. Two-way ANOVA for comparisons between the pcDNA3.1(+) empty vector group and other represented groups: p < 0.001 (***), p > 0.05 (ns). Values represent mean ± standard deviation.

**Figure S8.**
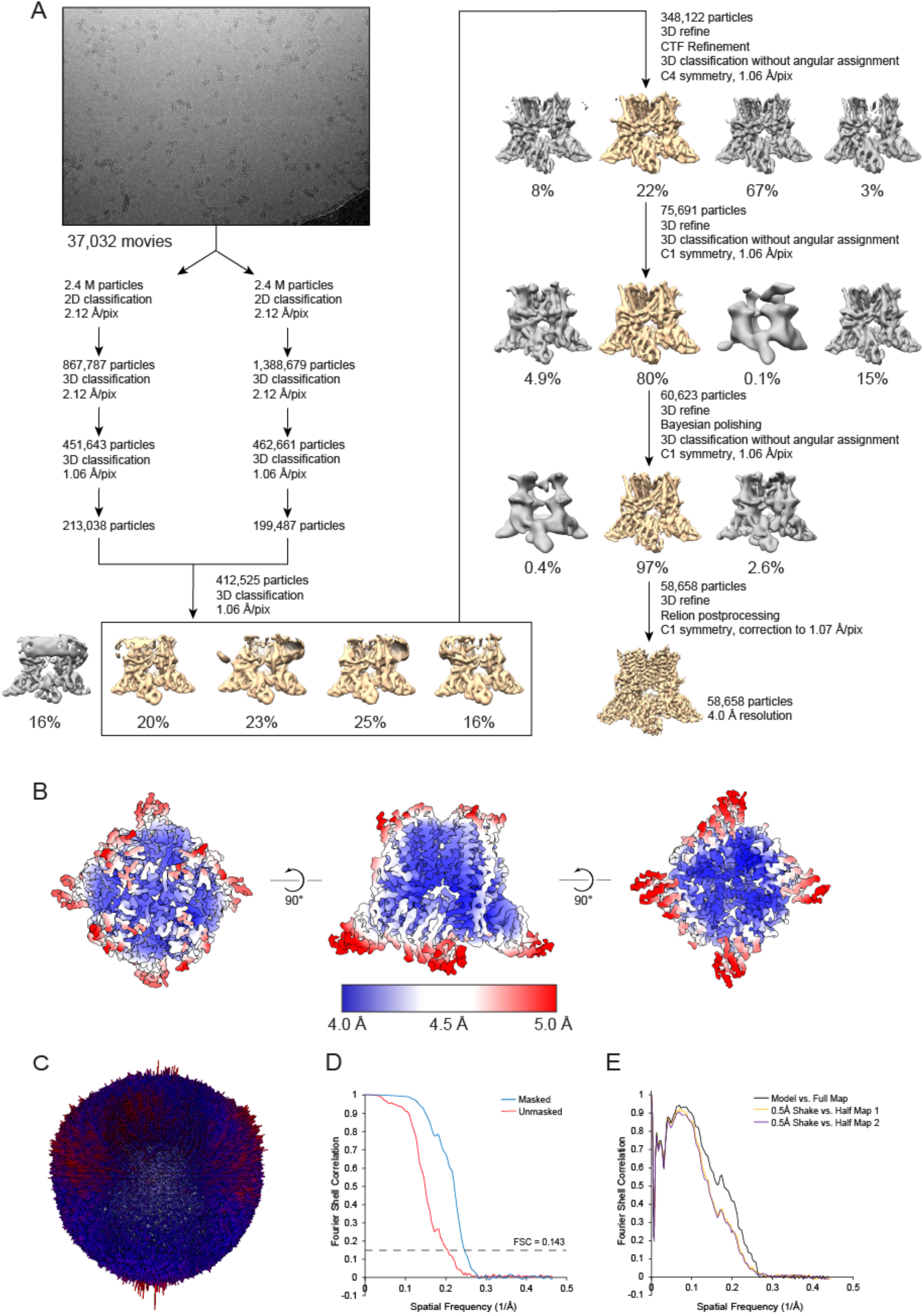
Overview of cryo-EM data from TRPV2_DOXO_. (A) Diagram of the data processing path from Relion. (B) Local resolution of the TRPV2_DOXO_ map. (C) Angular distribution of particles. FSC curves of the final map (D) and model validation (E).

**Figure S9.**
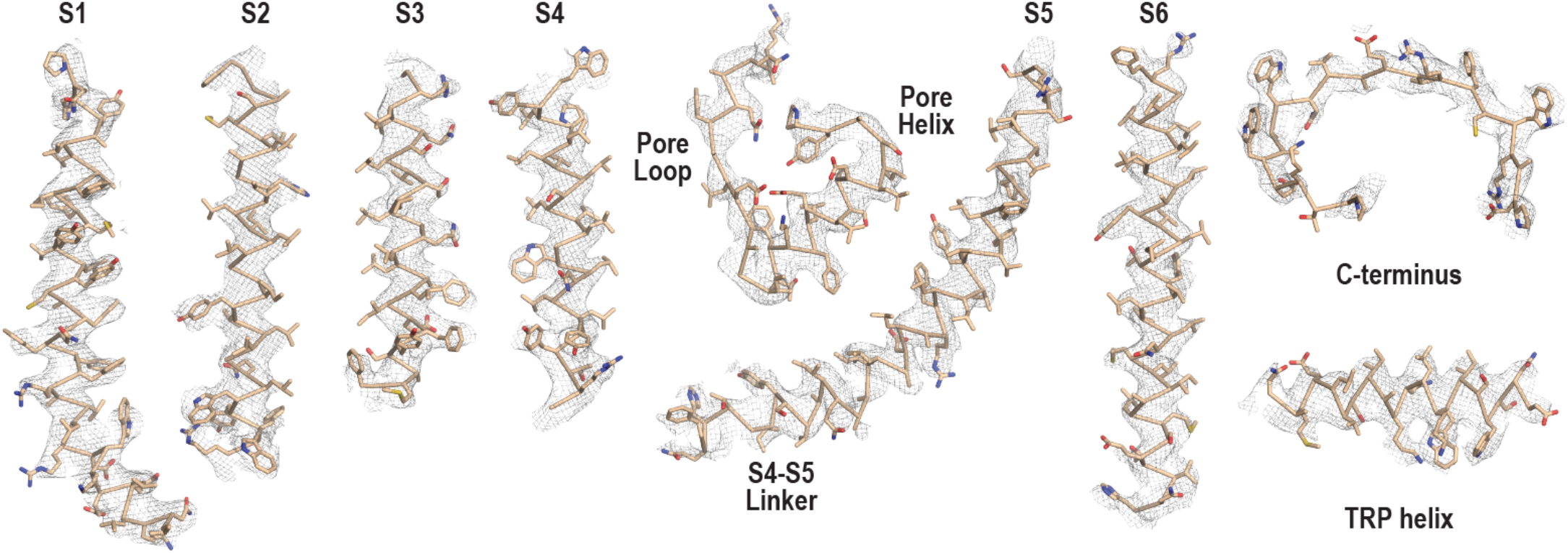
Representative densities of the TRPV2_DOXO_ cryo-EM map. Model is depicted as a ribbon with sidechains as sticks; densities are contoured at σ = 5.0.

**Figure S10.**
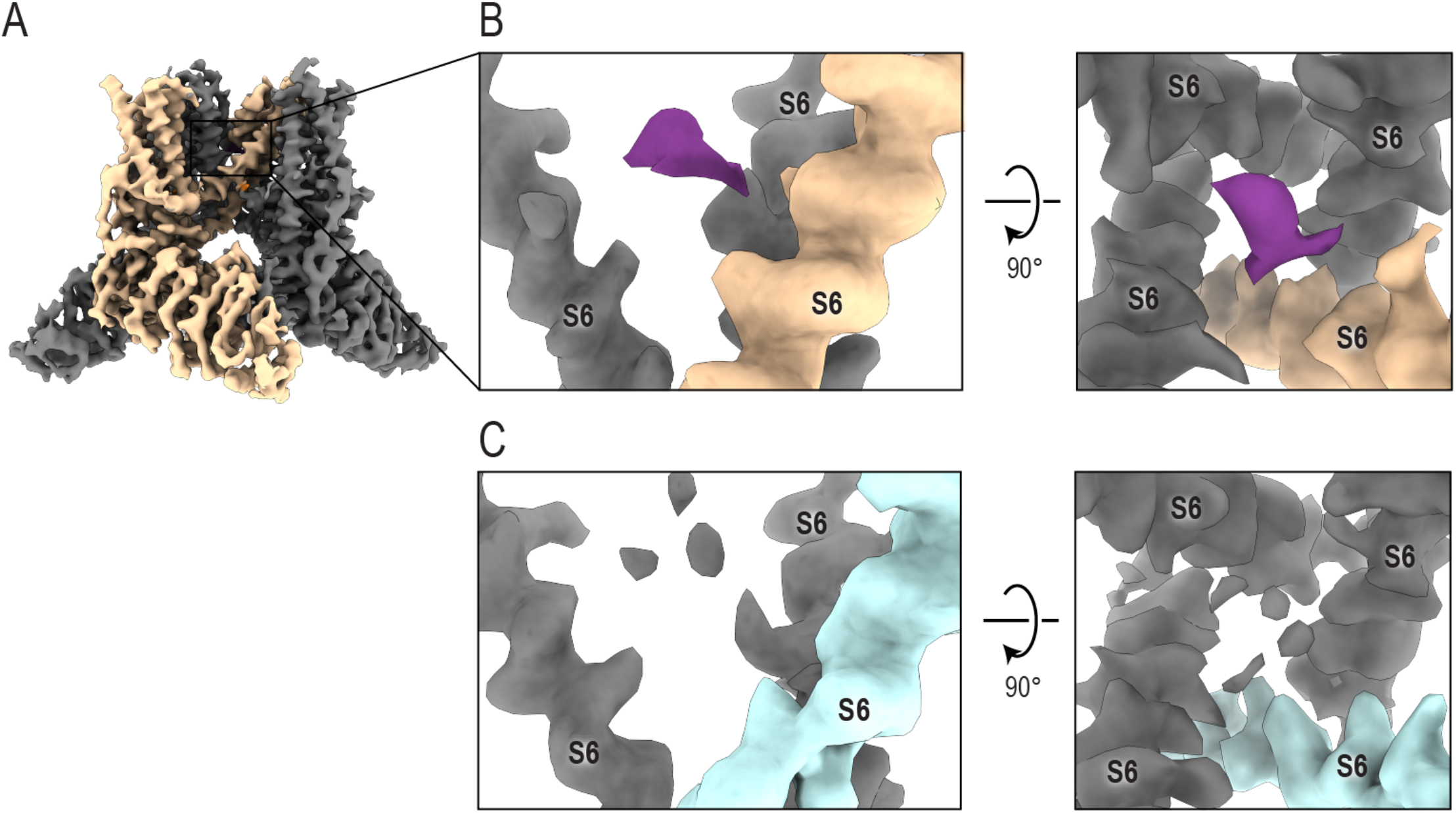
Cryo-EM density for DOXO in the TRPV2 pore. (A) Surface representation of the TRPV2_DOXO_ cryo-EM map. Detailed view of (B) the DOXO binding site and (C) the same site from the final particles of the TRPV2_2APB_ cryo-EM map refined without symmetry. TRPV2 is colored wheat (TRPV2_DOXO_) or pale cyan (TRPV2_2APB_); density attributed to DOXO is colored purple.

**Figure S11.**
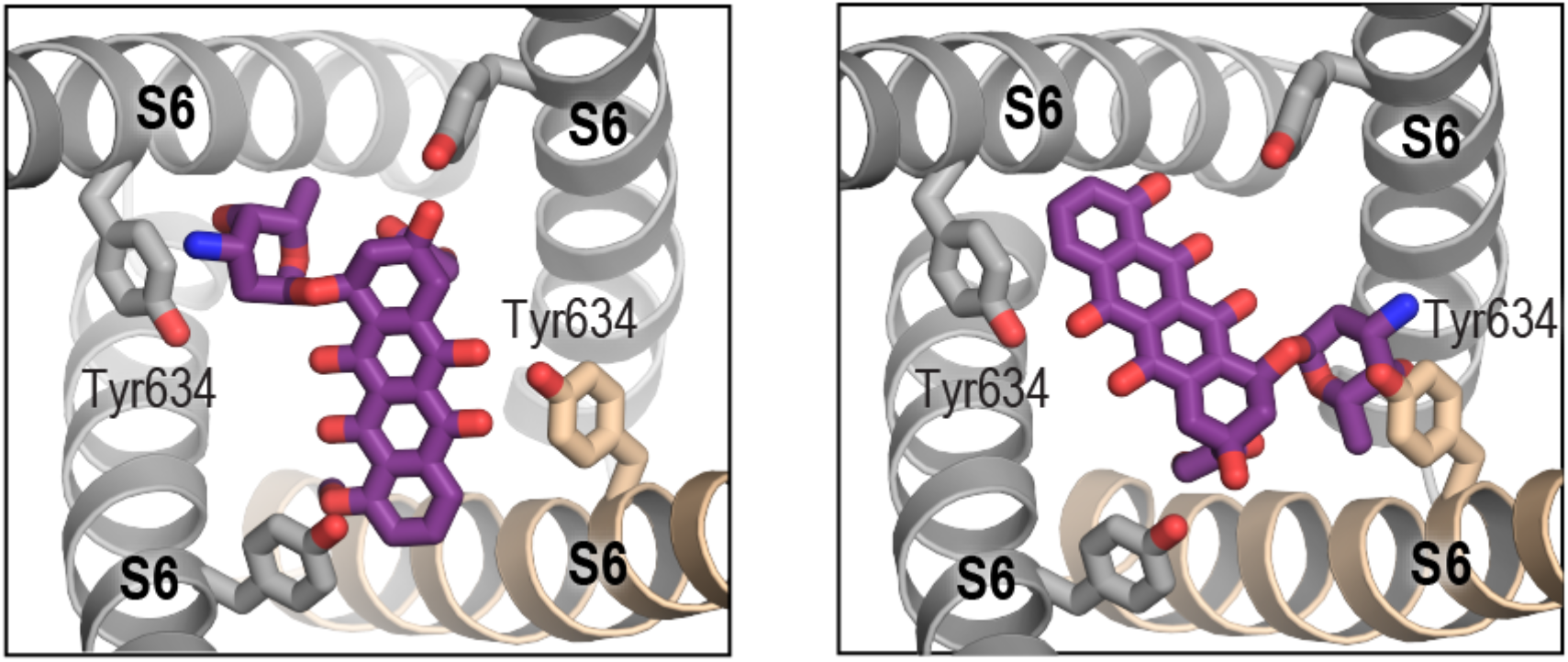
Two possible orientations of the DOXO molecule in the EM density.

**Figure S12.**
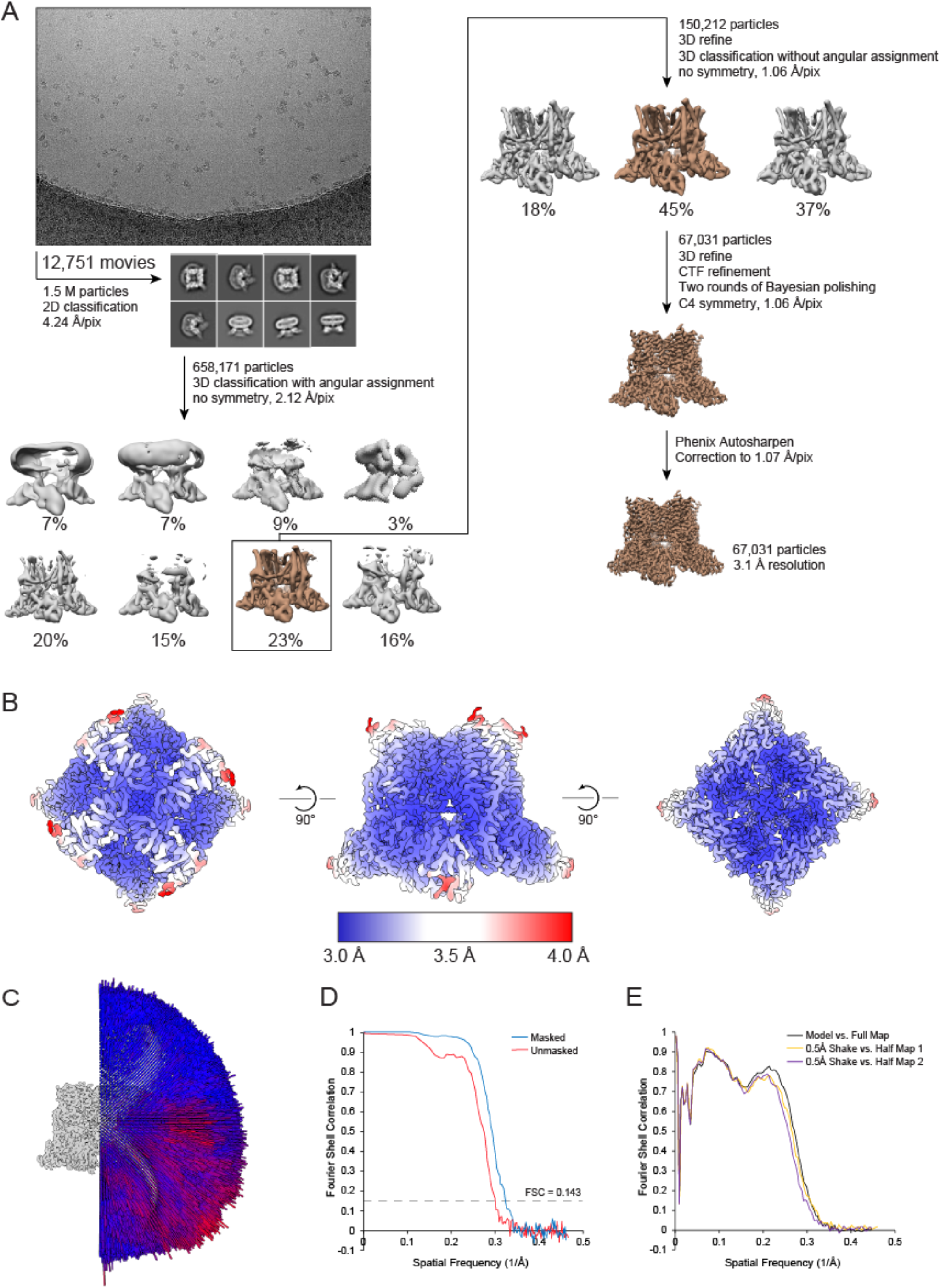
Overview of cryo-EM data from TRPV2_RR_. (A) Diagram of the data processing path from Relion. (B) Local resolution of the TRPV2_RR_ map. (C) Angular distribution of particles. FSC curves of the final map (D) and model validation (E).

**Figure S13.**
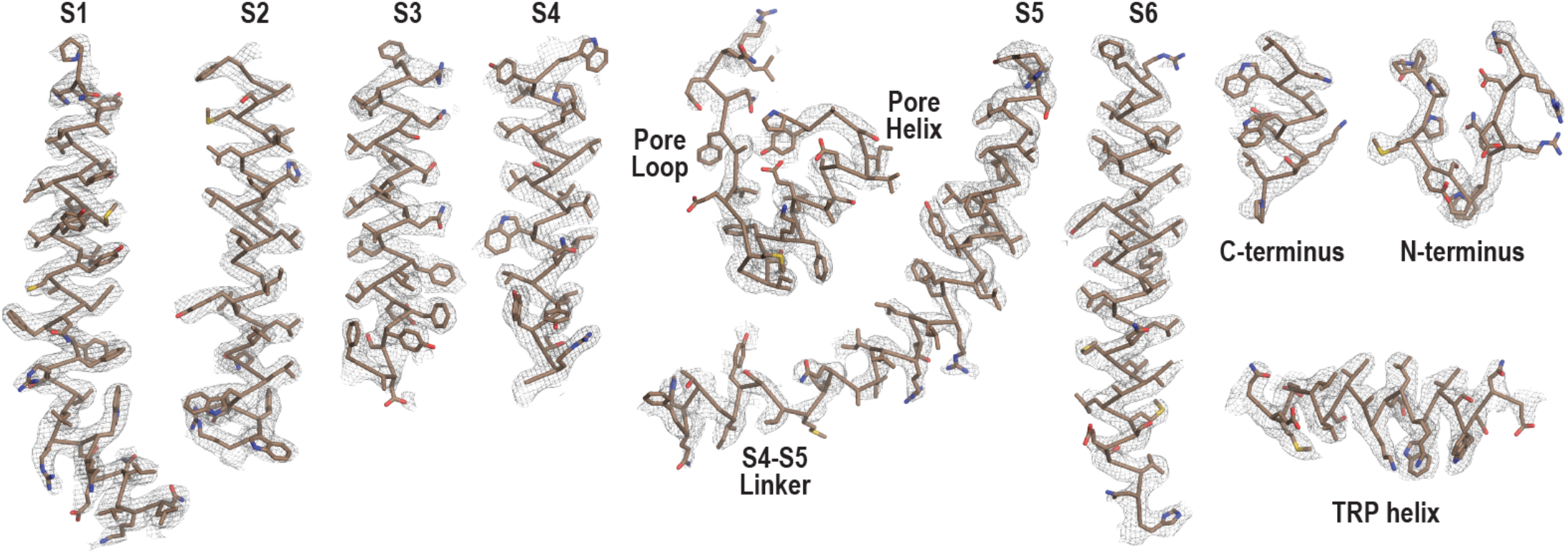
Representative densities of the TRPV2_RR_ cryo-EM map. Model is depicted as a ribbon with sidechains as sticks; densities are contoured at σ = 4.2.

**Figure S14.**
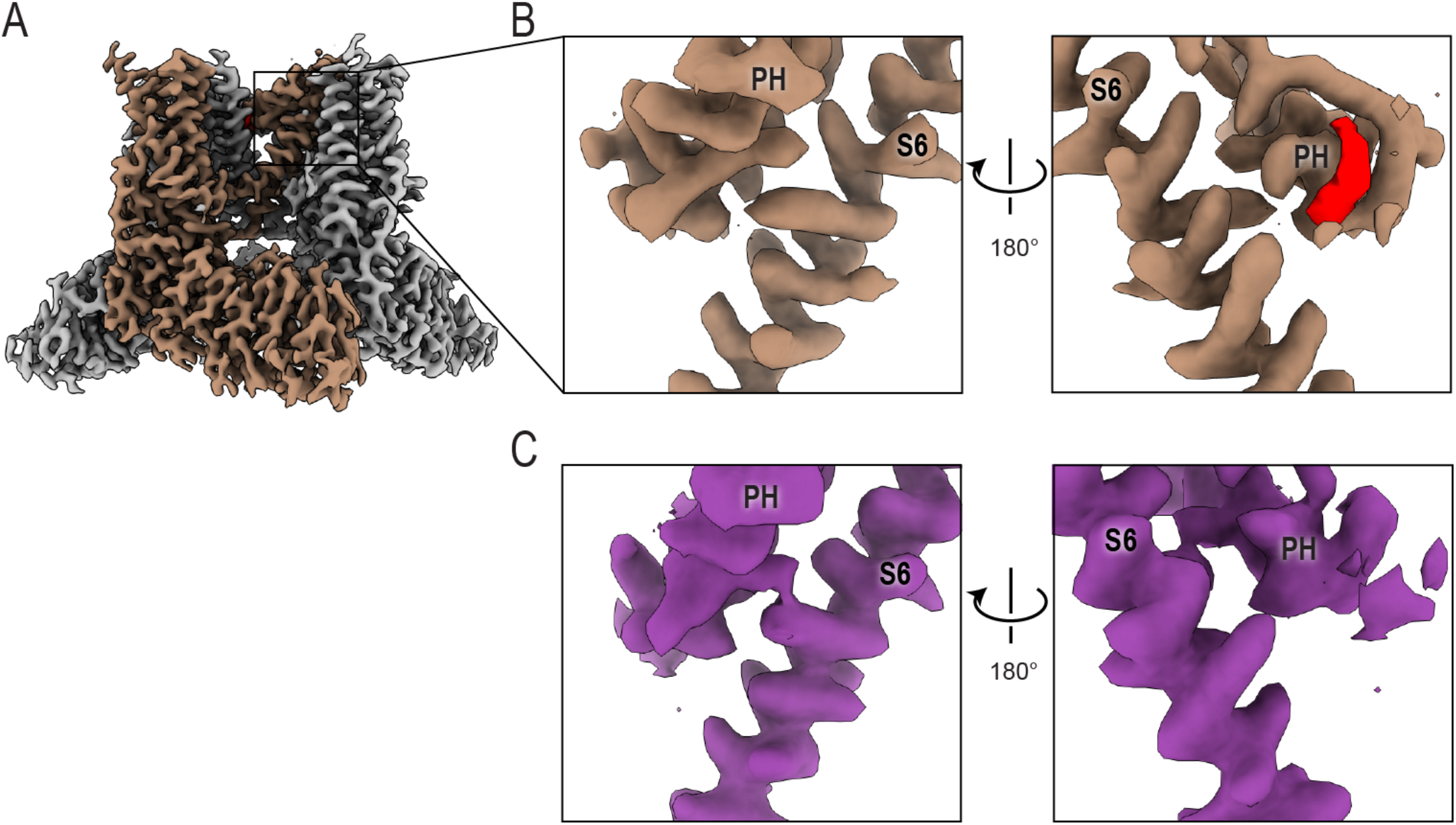
Cryo-EM density for RR at the selectivity filter. (A) Surface representation of the TRPV2_RR_ cryo-EM map. Detailed view of (B) the RR binding site and (C) the same site from TRPV_2APO_2_ (EMD-20678). TRPV2 is colored brown (TRPV2_RR_) or purple (TRPV_2APO_2_); density attributed to RR is colored red.

**Figure S15.**
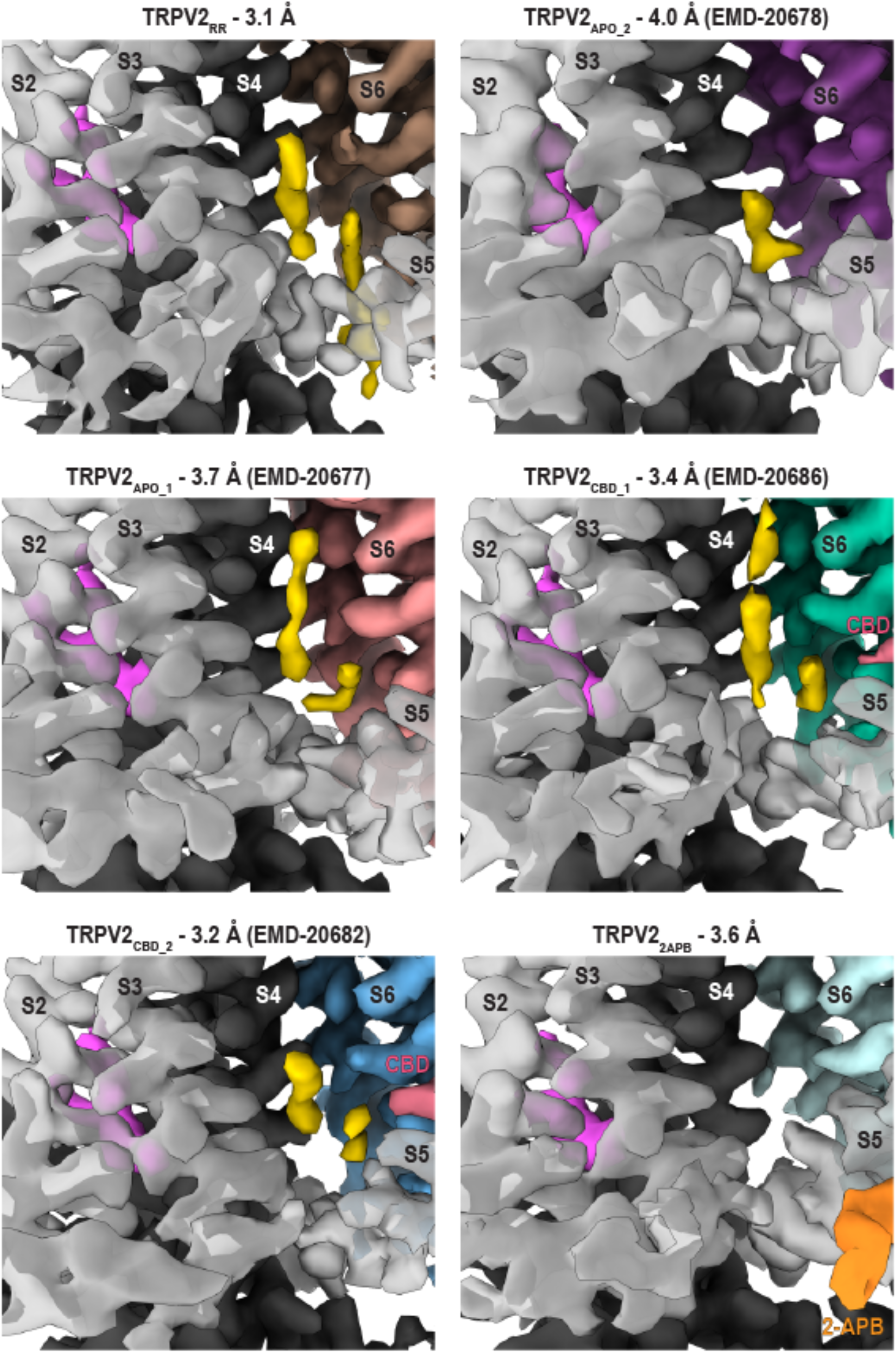
Cryo-EM density in the vanilloid pocket of TRPV2 structures in nanodiscs. Surface representations of the cryo-EM density of available TRPV2 structures in nanodiscs. Vanilloid lipid density is colored yellow, VSLD lipid density is colored magenta. Density for one monomer of each structure is colored grey, while the adjacent monomer is colored brown (TRPV2_RR_), purple (TRPV_2APO_2_, EMD-20678), salmon (TRPV_2APO_1_, EMD-20677), green (TRPV_2CBD_1_, EMD-20686), blue (TRPV_2CBD_2_, EMD-20682), or paly cyan (TRPV2_2APB_). Density for 2-APB is colored orange, density for CBD is colored pink.

**Figure S16.**
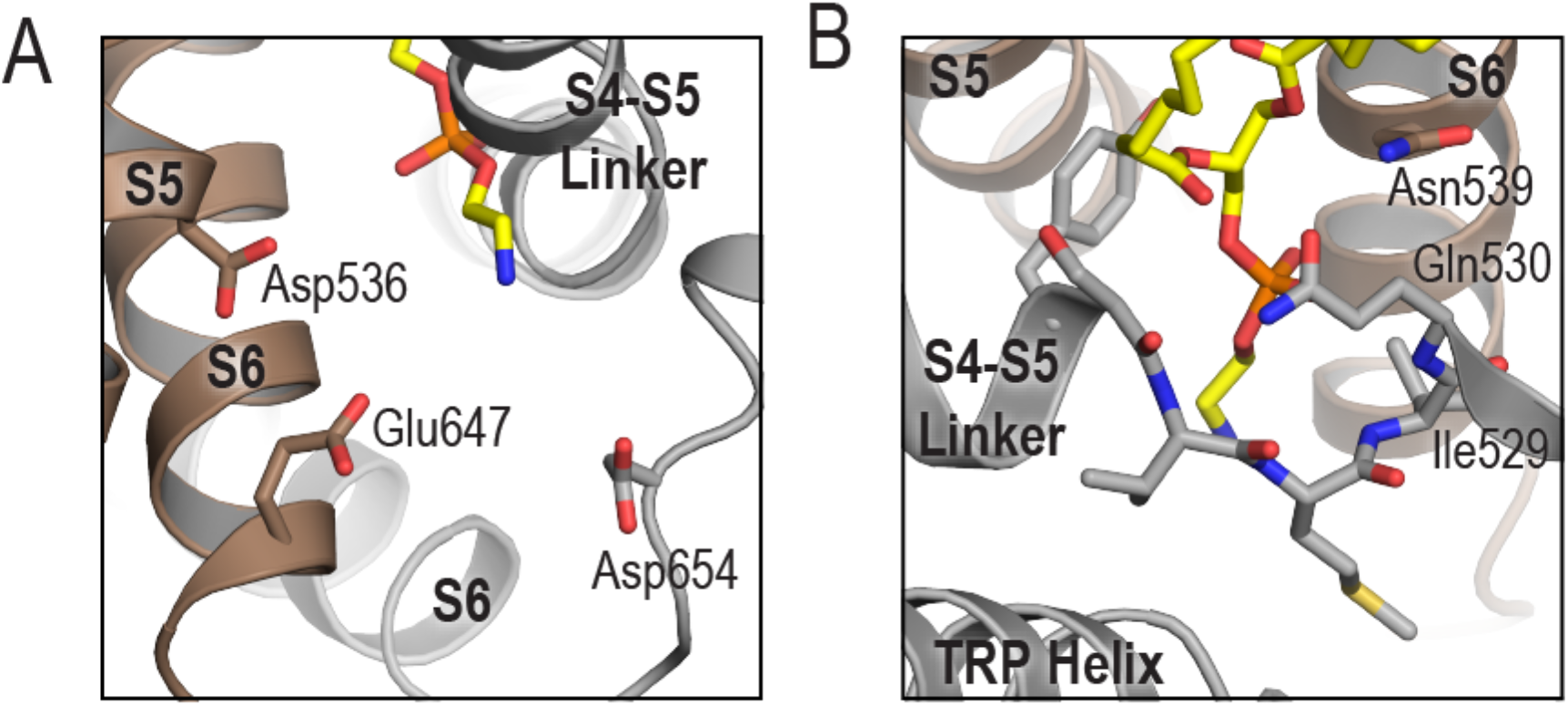
A docked lipid in the vanilloid pocket. Cartoon representation of the TRPV2_RR_ model with a phosphatidylethanolamine docked into the vanilloid pocket. (A) View of the acidic residues surrounding the proposed lipid headgroup. (B) View of the residues involved in stabilizing the proposed lipid in the vanilloid pocket. One monomer is colored brown, the adjacent monomer is colored grey, and the lipid is colored yellow. The lipid and residues of interest are depicted as sticks.

**Figure S17.**
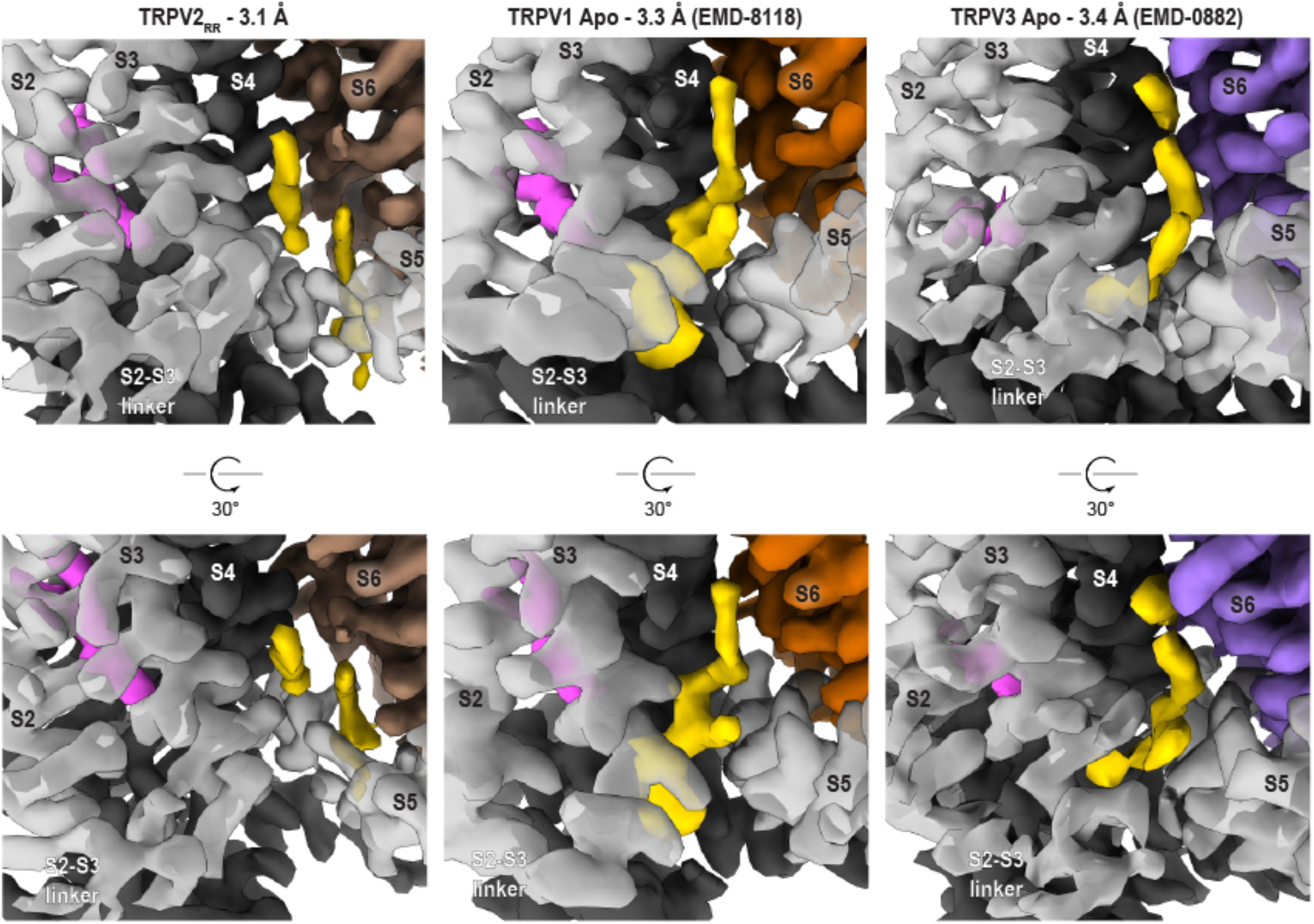
Lipids in the vanilloid pockets of TRPV1-3. Surface representation of cryo-EM density of TRPV_RR_, apo TRPV1 in nanodiscs (EMD-8118), and apo TRPV3 in nanodiscs (EMD-0882). Vanilloid lipid density is colored yellow, VSLD lipid density is colored magenta. Density for one monomer of each structure is colored grey, while the adjacent monomer is colored brown (TRPV2), orange (TRPV1), or purple (TRPV3).

